# Regenerative human liver organoids (HLOs) in a pillar/perfusion plate for hepatotoxicity assays

**DOI:** 10.1101/2024.03.25.586638

**Authors:** Sunil Shrestha, Prabha Acharya, Soo-Yeon Kang, Manav Goud Vanga, Vinod Kumar Reddy Lekkala, Jiafeng Liu, Yong Yang, Pranav Joshi, Moo-Yeal Lee

**Author notes:** Corresponding author: Moo-Yeal Lee. **Abbreviations:** LGR5, leucine-rich-repeat-containing G-protein-coupled receptor 5; Exp-HLOs, expandable human liver organoids; EM, expansion medium; Diff-HLOs, Differentiated human liver organoids; DM, differentiation medium; DILI, drug-induced liver injury.

## Abstract

Human liver organoids (HLOs) differentiated from embryonic stem cells (ESCs), induced pluripotent stem cells (iPSCs), and adult stem cells (ASCs) can recapitulate the structure and function of human fetal liver tissues, thus being considered as a promising tissue model for liver diseases and predictive compound screening. However, the adoption of HLOs in drug discovery faces several technical challenges, which include the lengthy differentiation process with multiple culture media leading to batch-to-batch variation, short-term maintenance of hepatic functions post-maturation, low assay throughput due to Matrigel dissociation and HLO transfer to a microtiter well plate, and insufficient maturity levels compared to primary hepatocytes. To address these issues, expandable HLOs (Exp-HLOs) derived from human iPSCs were generated by optimizing differentiation protocols, which were rapidly printed on a 144-pillar plate with sidewalls and slits (144PillarPlate) and dynamically cultured for up to 20 days into differentiated HLOs (Diff-HLOs) in a 144-perfusion plate with perfusion wells and reservoirs (144PerfusionPlate) for *in situ* organoid culture and analysis. The dynamically cultured Diff-HLOs exhibited greater maturity and reproducibility than those cultured statically, especially after a 10-day differentiation period. In addition, Diff-HLOs in the pillar/perfusion plate were tested with acetaminophen and troglitazone for 3 days to assess drug-induced liver injury (DILI) and then incubated in an expansion medium for 10 days to evaluate liver recovery from DILI. The assessment of liver regeneration post-injury is critical to understanding the mechanism of recovery and determining the threshold drug concentration beyond which there will be a sharp decrease in the liver’s regenerative capacity. We envision that bioprinted Diff-HLOs in the pillar/perfusion plate could be used for high-throughput screening (HTS) of hepatotoxic compounds due to the short-term differentiation of passage-able Exp-HLOs, stable hepatic function post-maturation, high reproducibility, and high throughput with capability of *in situ* organoid culture, testing, staining, imaging, and analysis.

**Graphical abstract:** The overall process of dynamic liver organoid culture and *in situ* analysis in the 144PillarPlate/144PerfusionPlate for high-throughput hepatotoxicity assays.

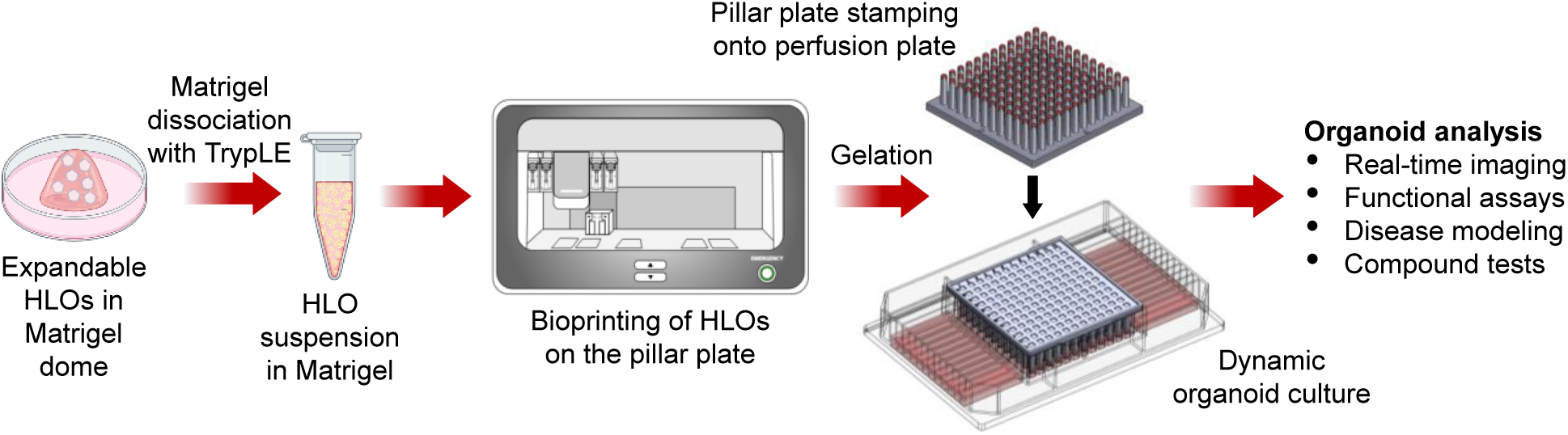

## Introduction

Sandwich-cultured primary human hepatocytes (PHHs) have been used widely as the gold standard of an *in vitro* liver model for several decades for disease modeling and hepatotoxicity testing in preclinical studies^1,2^. Nevertheless, PHHs are expensive and difficult to obtain in large quantities for high-throughput screening (HTS) of compounds^3^. When PHHs are maintained under standard *in vitro* cell culture conditions, they rapidly lose liver-specific functions over time^4^. Additionally, PHHs show high donor-to-donor variability in hepatic functions, leading to irreproducible results and significant lab-to-lab variability.

To address the limited availability of PHHs, human liver organoids (HLOs) differentiated from induced pluripotent stem cells (iPSCs) and adult stem cells (ASCs) have been extensively studied because of their cellular structure and function being strikingly similar to human fetal liver tissues^5^. In 2013 and 2015, Huch et al.^6,7^ generated biliary stem cell organoids from liver biopsy samples and demonstrated the generation of functional hepatocytes and cholangiocytes through further differentiation. In 2018, Hu et al.^8^ generated human hepatocyte organoids by expanding primary human hepatocytes. Nonetheless, HLOs derived from adult and pediatric liver biopsy samples had limited expansion capacity (up to 2 - 3 months), although fetal tissue-derived hepatocyte organoids did not show senescence. Additionally, liver tissue-derived HLOs have limited potential in modeling complicated liver diseases that require multiple hepatic cell types such as hepatocytes, stellate cells, Kupffer cells, and endothelial cells^9^. Furthermore, the variable efficiency of liver tissue-derived HLO generation raises concerns for patient-specific liver disease modeling and scale-up production of HLOs for compound screening.

On the other hand, HLOs derived from iPSCs demonstrated enhanced complexity and could potentially mimic the *in vivo* pathophysiology because iPSCs showed the ability to differentiate into multiple hepatic cell types. Several differentiation protocols have been developed to generate HLOs using iPSCs alone^10–15^ or iPSCs with other cell types^16–18^. Among hepatic differentiation protocols, Ouchi et al.^13^ pioneered the generation of iPSC-derived multicellular HLOs consisting of parenchymal and non-parenchymal liver cell types for modeling steatohepatitis. They showcased inflammatory and fibrotic responses of steatohepatitis by 5-day treatment with fatty acids. By using the same differentiation protocol, Shinozawa et al.^15^ demonstrated bile acid uptake by HLOs, a critical feature of functional bile canaliculi in the liver and performed large-scale compound screening for cholestatic liver injury using live HLO imaging for bile acid uptake and viability. This differentiation of iPSCs into multicellular HLOs is robust, but it requires month-long cell differentiation with several cell culture media. Additionally, mature HLOs gradually lose their hepatic functions over time, which could be a critical issue for reproducible organoid-based assays. For scale-up production of HLOs, foregut cells are embedded in Matrigel domes, which need to be physically or enzymatically dissociated to isolate and harvest mature HLOs. This process is labor-intensive and could lead to the loss of HLOs, damage to HLO structure and function, and batch-to-batch variation in their size.

Harvested HLOs, 200 – 300 µm in diameter, need to be dispensed in a microtiter well plate for organoid-based assays^15^, which could require an expensive liquid handling machine for high accuracy.

To resolve these issues, expandable human liver organoids (Exp-HLOs) were generated in this work by differentiating day 15 early hepatic progenitor cells derived from iPSCs, which can be passaged and cryopreserved for the scale-up production of HLOs. Subsequently, functional differentiated human liver organoids (Diff-HLOs) were generated by differentiating the Exp-HLOs. To enhance the maturity of Diff- HLOs, we optimized the differentiation medium and culture duration and incorporated a pillar/perfusion plate for dynamic organoid culture. Additionally, this platform supported the uniform generation of Diff- HLOs as well as high-throughput assessment of drug-induced liver injury (DILI) and the recovery of the liver from DILI using Diff-HLOs. Thus, we envision that our approach could be a valuable addition to current *in vitro* hepatotoxicity assessment which mainly focuses on live-dead viability of primary hepatocytes.

## Materials and Methods

### Fabrication of 144PillarPlate and 144PerfusionPlate

The 144PillarPlate, the 144PerfusionPlate, and the 384DeepWellPlate (**Figure 4 and Supplementary Figure 1**) were manufactured by injection molding of polystyrene (Bioprinting Laboratories Inc., Dallas, TX, USA). The 144PilllarPlate contains a 12 x 12 array of pillars (4.5 mm pillar-to-pillar distance, 11.6 mm pillar height, and 2.5 mm outer and 1.5 mm inner diameter of pillars), with each pillar allowing to load up to 6 µL of cells suspended in hydrogel. The 144PerfusionPlate has a 12 x 12 array of perfusion wells (3.4 mm width, 3.4 mm length, 11.9 mm depth, and 4.5 mm well-to-well distance) with a 2 x 12 array of reservoirs (3.6 mm width, 30.3 mm length, 15.4 mm depth, and 4.5 mm reservoir-to-reservoir distance), all connected by microchannels. Thus, the 144PerfusionPlate contains 12 fluidic channels, each channel consisting of 2 reservoirs (upper and lower) on both sides and 12 perfusion wells in a row connected by microchannels, allowing up to 2,000 µL of cell growth medium for dynamic organoid culture. The 384DeepWellPlate has a 16 x 24 array of deep wells (3.5 mm width, 3.5 mm length, 14.7 mm depth, and 4.5 mm well-to-well distance), with each deep well allowing up to 80 µL of a cell growth medium for static organoid culture. As in our previous studies^19,20^, flow simulation in the perfusion plate was conducted using the SolidWorks software package (SolidWorks Research Standard and Simulation Premium 2022, MA, USA). The pillar plate with cells encapsulated in hydrogels can be sandwiched onto the perfusion plate or the deep well plate for dynamic and static culture of human organoids as well as *in situ* compound testing and organoid imaging.

### iPSCs maintenance and differentiation into expandable human liver organoids (Exp-HLOs)

EDi029-A, a male human iPSC line (Cedar Sinai Biomanufacturing Center, USA), was maintained on growth factor-reduced Matrigel (Corning; 354230) coated dishes under mTeSR^TM^ Plus medium (StemCell Technologies; 100-0276) and passaged using the StemPro^TM^ EZPassage^TM^ passaging tool (ThermoFisher; 23181-010) or ReLeSR^TM^ human PSC selection and passaging reagent (StemCell Technologies; 05872). The differentiation of iPSCs into foregut cells was performed using previously published protocols^13,21^. Briefly, at 70 - 80% confluency, iPSCs were harvested using Accutase (Gibco; A1110501), seeded on iMatrix-511 silk (Elixirgen Scientific; NI511) coated 6-well plate at a cell density of 1.3 × 10^6^, and cultured using mTESR^TM^ Plus medium supplemented with 10 µM Y27632 Rho-kinase (ROCK) inhibitor (Tocris; 1254). After 24 hours of culture, the iPSCs were differentiated into definitive endoderm using RPMI 1640 (Gibco; 22400089) supplemented with 50 ng/mL bone morphogenetic protein 4 (BMP4; Tocris; 314-BP) and 100 ng/mL activin A (Tocris; 338-AC) on day 1, 100 ng/mL Activin A and 0.2% (v/v) knockout serum replacement (KSR; Gibco; 10828010) on day 2, and 100 ng/mL Activin A and 2% (v/v) KSR on day 3. This was followed by foregut cell differentiation from day 4 - 6 using advanced DMEM/F12 (Gibco; 12634) with 2% (v/v) B27 (Gibco; 17504), 1% (v/v) N2 (Gibco; 17502), 10 mM HEPES (Gibco; 15630), 1% (v/v) penicillin/streptomycin (Gibco; 15140), and 1% (v/v) GlutaMAX^TM^ (Gibco; 35050) supplemented with 500 ng/mL fibroblast growth factor (FGF4; Peprotech; 100-31) and 3 µM CHIR99021 (R&D Systems; 4423). On day 7, foregut cells were dissociated using Accutase, resuspended in Matrigel (Corning; 356237) at a density of 750 cells/µL, and dispensed in a 24-well plate to form 50 µL Matrigel domes for culture, or cryopreserved using CryoStor^®^ CS10 (StemCell Technologies; 07959) for later use. After gelation of Matrigel at 37°C for 10 - 12 minutes, foregut cells in the 24-well plate were cultured in DMEM base medium supplemented with 5 ng/mL recombinant human FGF basic/FGF2/bFGF (R&D systems; 233-FB), 10 ng/mL recombinant human VEGF-165 (Gibco; PHC9391), 20 ng/mL recombinant human EGF (R&D system; 236-EG), 3 µM CHIR99021, 0.5 µM A 83-01 (R&D Systems; 2939), 50 µg/mL L-ascorbic acid (Sigma; A4544), and the CEPT cocktail consisting of 50 nM chroman 1 (R&D systems; 7163), 5 µM emricasan (Selleckchem; S7775), 0.1% (v/v) polyamine supplement (Sigma; P8482), and 0.7 µM trans-ISRIB (R&D systems; 5284). The medium volume used in the 24-well plate was 1 mL per well. The differentiation medium was changed every other day. On day 11, the differentiation medium was changed to DMEM base medium supplemented with 2 µM retinoic acid (Sigma; R2625) with medium changes every other day.

Starting from day 15, early hepatic progenitor cells was differentiated into expandable human liver organoids (Exp-HLOs) in an expansion medium (EM) consisting of advanced DMEM/F12 supplemented with 2% (v/v) B27 supplement (without vitamin A), 1% (v/v) N2 supplement, 10 mM HEPES, 1% (v/v) GlutaMAX, 1% (v/v) penicillin/streptomycin, 1 mM N-acetyl-L-cysteine (Sigma; A9165), 10 mM nicotinamide (Sigma; N0636), 10 nM recombinant human (Leu^15^)-gastrin I (Sigma; G9145), 50 ng/mL recombinant human EGF, 500 ng/mL R-Spondin1 (Rspo1) (R&D systems; 4645-RS/CF), 100 ng/mL recombinant human FGF10 (Peprotech; 100-26), 25 ng/mL recombinant human HGF (Peprotech; 100-39), 10 μM forskolin (R&D Systems; 1099), and 5 μM A83-01. The Exp-HLOs were cultured for 1 - 2 weeks until these organoids reached a size of approximately 250 µm or larger. The Exp-HLOs were enzymatically dissociated into small clumps by treatment with 1x TrypLE™ Express Enzyme (ThermoFisher; 12605010) for 5 minutes at 37°C in the CO_2_ incubator. The dissociated cell clumps were either cryopreserved using Stem-Cellbanker DMSO free - GMP grade (Amsbio; 13926) for later use or mixed with undiluted Matrigel (Corning; 356237) at the ratio of 1:10 for expansion. When cell clumps were thawed and cultured post-cryopreservation, the EM was supplemented with the cryoprotectant CEPT cocktail for the first 4 days of culture.

### Differentiation of passaged Exp-HLOs into Diff-HLOs

The dissociated cell clumps from Exp-HLOs were mixed with undiluted Matrigel (Corning; 356237) and dispensed at 50 μL Matrigel dome/well in a 24-well plate for differentiation into Diff-HLOs. The cell clumps in Matrigel dome were cultured in the EM supplemented with 25 ng/mL human active BMP7 recombinant protein (Gibco; PHC7204) initially for 3 - 6 days to regenerate and form Exp-HLOs using BMP7. After forming Exp-HLOs approximately 200 μm in diameter, the organoids were cultured in a differentiation medium (DM) consisting of advanced DMEM/F12 supplemented with 1% (v/v) B27, 1% (v/v) N2, 10 mM HEPES, 1% (v/v) penicillin/streptomycin, and 1% (v/v) GlutaMAX, 10 nM recombinant human (Leu^15^)-gastrin I, 100 ng/mL recombinant human FGF19 (Peprotech; 100-32), 25 ng/mL recombinant human hepatocyte growth factor (HGF; Peprotech; 100-39), 500 nM A8301 (R&D Systems; 2939), 10 μM notch inhibitor DAPT (Sigma; D5942), 25 ng/mL human active BMP7 recombinant protein, and 3 μM dexamethasone (Dex; Sigma; D4902) for up to 20 days with medium changes every other day. Later, 20 ng/mL human oncostatin M (OSM; Peprotech; 300-10) was also added in the DM to further enhance the maturity and functionality of Diff-HLOs.

### Microarray bioprinting of cell clumps in Matrigel on the pillar plate

The cell clumps obtained from enzymatic dissociation of Exp-HLOs were mixed with undiluted Matrigel and printed on the 144PillarPlate at 4 µL cell clumps in Matrigel per pillar using ASFA^TM^ Spotter V6 (MBD Co., Ltd., South Korea). After Matrigel gelation at 37°C for 10 - 12 minutes, the 144PillarPlate with cell clumps was sandwiched onto a complementary 384DeepWellPlate containing 80 µL of EM+BMP7 medium in each well for static HLO culture or coupled with the 144PerfusionPlate containing 1300 µL of EM+BMP7 medium per fluidic channel for dynamic HLO culture.

### Gene expression analysis via RT-qPCR

HLOs in Matrigel were either collected manually by pipetting in cold DPBS without calcium and magnesium (dPBS^-/-^) or isolated from Matrigel using Cultrex organoid harvesting solution (R&D Systems; 3700-100-01) according to the manufacturer’s recommended protocol, which allows non-enzymatic depolymerization of Matrigel. In the case of HLOs on the pillar plate, the pillar plate was sandwiched onto the deep well plate containing 80 µL of Cultrex organoid harvesting solution. The sandwiched plates were incubated for 30 minutes at 4°C and then centrifuged at 100 rcf for 10 – 20 minutes to detach the organoids. Total RNA was isolated from harvested cells by using the RNeasy Plus Mini Kit (Qiagen; 74134) following the manufacturer’s recommended protocol. cDNA was synthesized from 1 µg of RNA using the high-capacity cDNA reverse transcription kit (Applied Biosystems; 4368814). Real-time PCR was performed using PowerTrack^TM^ SYBR green master mix (Applied Biosystems; A46110) and forward/reverse primers from IDT Technology in the QuantStudio™ 5 Real-Time PCR System (Applied Biosystems; A28574). The cycle was run 40 times denaturation at 95°C for 30 sec, annealing at 58 - 62°C (depending on primer pair) for 45 sec, and extension at 72°C for 30 sec. The primers used are listed in **Supplementary Table 1**. The expression level of target genes was normalized to that of the housekeeping gene, glyceraldehyde 3- phosphate dehydrogenase (*GAPDH*).

### Whole mount immunofluorescence staining

Immunofluorescence staining was performed either by harvesting HLOs from Matrigel domes in the 24- well plate or with HLOs *in situ* on the pillar plate. In the case of Matrigel dome culture, Matrigel domes containing organoids were collected in cold dPBS^-/-^ through pipetting into a 1.5 mL Eppendorf tube and then centrifuged at 300 x g for 4 minutes to isolate HLOs. The HLOs were fixed using 4% (w/v) paraformaldehyde (PFA; ThermoFisher Scientific; J19943K2) for 2 hours at room temperature while gently rocking. The fixed HLOs were washed with dPBS^-/-^ containing 0.1% (w/v) sodium borohydride twice for 15 minutes to reduce background due to free aldehyde. After washing, the HLOs were permeabilized with 500 µL of 0.5% (v/v) Triton X-100 (Sigma; T8787) in dPBS^-/-^ (i.e., permeabilization buffer) for 15 - 20 minutes twice at room temperature with gentle rocking. After permeabilization, the HLOs were exposed to 500 µL of 5% (v/v) normal donkey serum (NDS) in the permeabilization buffer (i.e., blocking buffer) for 1 hour at room temperature or overnight at 4°C with gentle rocking to prevent non-specific binding. For primary antibody staining, the HLOs were treated with 250 µL of 5 µg/mL primary antibody diluted in the blocking buffer for overnight at 4°C with gentle rocking. The HLOs were rinsed with 1 mL of the blocking buffer thrice for 30 minutes each at room temperature with gentle rocking to prevent non-specific binding. For secondary antibody staining, the HLOs were exposed to 500 µL of 5 µg/mL fluorophore-conjugated secondary antibody in the blocking buffer for 2 - 4 hours at room temperature with gentle rocking. The HLOs were stained with 500 µL of 0.5 µg/mL DAPI solution (ThermoFisher Scientific; 62248) in 1x dPBS^-^

^/-^ for 30 minutes at room temperature with gentle rocking. The HLOs were further washed with 1 mL of dPBS^-/-^ twice to ensure the complete removal of unbound secondary antibody. Finally, the HLOs were transferred to a microscope cover glass (Fisher Scientific; 22266882) and treated with 25 μL of Visikol^®^ Histo-M™ (Visikol; HM-30) to clear the organoids which also works as a mounting solution. The HLOs on the cover glass slide were covered by another cover glass from the top and imaged using a Zeiss LSM 710 confocal scanning microscope. For the HLOs on the pillar plate, all the immunofluorescence staining steps were performed by sandwiching the pillar plate with a 384DeepWellPlate containing 80 µL of respective solutions and incubating the sandwiched plates under the same conditions mentioned above. After the final wash with 80 μL dPBS^-/-^ to remove unbound secondary antibody, the HLOs were treated with 35 µL of Visikol^®^ Histo-M™ in a regular 384-well plate (ThermoFisher Scientific; 242757) for 1 hour at room temperature. At the time of imaging, the pillar plate containing stained HLOs was placed on the microscope cover glass. The specific names of primary and secondary antibodies used are listed in **Supplementary Table 2 and 3**, respectively.

### Cell cytometry analysis

Exp-HLOs in Matrigel were collected in cold dPBS^-/-^ and dissociated into single cells using Accutase for 5 - 8 minutes at 37° C in a CO_2_ incubator. The dissociated cells were collected in a 15 mL tube and fixed using 4% PFA for 15 minutes at room temperature. Subsequently, the cells were permeabilized with 0.25% Triton X-100 in dPBS^-/-^ for 15 minutes and blocked using 5% NDS in dPBS^-/-^ containing 0.1% Triton X- 100 for 30 minutes at room temperature. The cells were then incubated with 2.5 µg/mL of EpCAM (SantaCruz; sc-25308) for 1 hour at room temperature, washed with the blocking solution thrice, and stained with 5 µg/mL of donkey anti-mouse IgG (H+L) highly cross-adsorbed secondary antibody, Alexa Fluor™ 488 (Invitrogen; A-21202) for 30 minutes at room temperature. After staining, the cells were washed with dPBS^-/-^ three times before flow cytometry analysis. UltraComp eBeads™ compensation beads (Fisher Scientific; 01-2222-41) stained with the same secondary antibody were used as a positive control for analysis. The assessment was performed using the Cytek Aurora spectral flow cytometer (Cytek^®^ Biosciences) and analyzed with FlowJo 10.9.0 (Becton Dickinson & Company).

### Measurement of cell viability

The viability of Exp-HLOs was analyzed using CellTiter-Glo^®^ 3D cell viability assay kit (Promega; G9681) following the manufacturer’s recommended protocol. Briefly, the pillar plate with HLOs was sandwiched with an opaque white 384-well plate containing a mixture of 30 µL of the CellTiter-Glo^®^ reagent and 10 µL of the cell culture medium in each well to measure cellular adenosine triphosphate (ATP) levels. To induce cell lysis, the sandwiched pillar/well plates were placed on an orbital shaker for 1 hour at room temperature. After cell lysis, the pillar plate was detached, and the lysis solution in the opaque white 384- well plate was left for 15 minutes at room temperature for stabilization. The luminescence signals were recorded using a microtiter well plate reader (BioTek^®^ Cytation 5).

The viability of Exp-HLOs in a 24-well plate was also analyzed by using the Cell Counting Kit-8 (CCK- 8; Dojindo; CK04). The CCK-8 kit is non-toxic to organoids, thus allowing to monitor cell proliferation continuously. Briefly, the Exp-HLOs in the 24-well plate were treated with the CCK-8 solution diluted 10- fold in the EM without Rspo1 by mixing 30 μL of the CCK-8 solution with 270 μL of the EM without Rspo1 (EM-Rspo1) in each well. After 2.5 hours of treatment in the CO_2_ incubator, 50 μL of the supernatant was dispensed in triplicate in the opaque white 384-well plate. Subsequently, 500 μL of fresh EM-Rspo1 was dispensed in the 24-well plate to continue culturing the organoids. The supernatant was left for 10 minutes at room temperature for stabilization, and the absorbance was measured at 450 nm by using the BioTek^®^ Cytation 5 plate reader.

### Measurement of CYP3A4 expression

The expression level of CYP3A4 was analyzed using the P450-Glo^TM^ CYP3A4 assay kit (Promega; V9001) following the manufacturer’s recommended protocol. Rifampicin (Sigma; R3501) was used as an inducer of the *CYP3A4* gene at the concentration of 25 µM. Briefly, on day 20, Diff-HLOs on the pillar plate were treated with rifampicin for 3 days with daily medium changes. After treatment, HLOs were incubated with luciferin IPA-substrate at a final concentration of 3 µM diluted in advanced DMEM/F12 for overnight at 37°C in the CO_2_ incubator. Following the overnight incubation, 25 µL of the culture medium from each well of the 384DeepWellPlate was transferred to an opaque white 384-well plate at room temperature, and 25 µL of luciferin detection reagent was added in each well to initiate a luminescent reaction. After 20 minutes of incubation at room temperature, luminescence was measured by using the BioTek^®^ Cytation 5 plate reader.

### Measurement of bile acid transport with cholyl-lysyl-fluorescein (CLF)

To analyze the function of bile acid transport in HLOs, the Diff-HLOs were washed with dPBS^-/-^ and incubated in the DM containing 5 µM cholyl-lysyl-fluorescein (Corning; 451041). After overnight treatment at 37°C in the CO_2_ incubator, the Diff-HLOs were rinsed with dPBS^-/-^ thrice and replaced with the DM without CLF. The Diff-HLOs were then imaged by using an automated fluorescence microscope (Keyence; BZ-X800E) equipped with a 20x objective lens. CLF is a fluorescein-labeled bile acid that can be transported into bile canaliculi by the bile salt efflux pump (BSEP).

### Testing model compounds with Diff-HLOs on the pillar plate

Day 20 Diff-HLOs on the pillar plate were exposed to varying concentrations of two hepatotoxic drugs, acetaminophen (Sigma; A5000) and troglitazone (Sigma; T2573). The highest dosage tested was 10,000 μM for acetaminophen and 500 μM for troglitazone. Briefly, 4-fold serial dilutions of the highest dose of the drugs were performed in DMSO in a 384-well plate. Five dosages and one solvent-alone control (DMSO control) were prepared for each drug. The drug stock solutions in DMSO in the 384-well plate were 200- fold diluted with the DM and then dispensed in the 144PerfusionPlate (duplicates per dose) so that the final DMSO content was equal to 0.5% (v/v). The pillar plate with day 20 Diff-HLOs was then sandwiched with the 144PerfusionPlate containing the serially diluted drugs and incubated in the 5% CO_2_ incubator at 37°C for 3 days. The viability of HLOs was assessed with the CellTiter-Glo^®^ 3D cell viability assay kit (Promega), and luminescence was measured using the BioTek^®^ Cytation 5 plate reader. Dose-response curves were generated using the luminescence values at varying dosages. To assess the recovery of the Diff-HLOs after drug treatment, the pillar plate with day 20 Diff-HLOs, exposed to the concentration of the drugs near to their IC_50_ value, was sandwiched with the 144PerfusionPlate containing the EM and incubated in the 5% CO_2_ incubator at 37°C for 10 days.

### Calculation of the IC_50_ value

Since the background luminescence of completely dead cells (following treatment with 70% methanol for 1 hour) was negligible due to background subtraction, the percentage of live Diff-HLOs was calculated using the following equation:

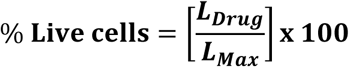

where L_Drug_ is the luminescence intensity of Diff-HLOs exposed to the drugs and L_Max_ is the luminescence intensity of fully viable Diff-HLOs (control).

To generate a conventional sigmoidal dose-response curve with response values normalized to span the range from 0% to 100% plotted against the logarithm of test concentration, we normalized the luminescence intensities of all Diff-HLO spots with the luminescence intensity of a 100% live Diff-HLO spot (Diff-HLOs incubated with no compound). We then converted the test drug concentrations to their respective logarithms using GraphPad Prism 9.3.1 (GraphPad Software, Inc., CA, USA). The sigmoidal dose-response curve (variable slope) and IC_50_ value (i.e., the concentration of drug where 50% of Diff-HLO viability is inhibited) were obtained using the following equation:

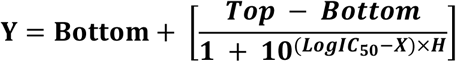

where IC_50_ is the midpoint of the curve, H is the hill slope, X is the logarithm of test concentration, and Y is the response (% live cells), starting from the top plateau (Top) of the sigmoidal curve to the bottom plateau (Bottom).

### Statistical analysis

The statistical analysis was performed using GraphPad Prism 9.3.1. All the data were expressed as mean ± SD, with sample sizes specified in the figure legends where ‘n’ represents biological replicates. Student’s t-test was used for comparison between two groups, whereas one-way ANOVA was used for comparison among multiple groups. The statistically significant difference between the control and test groups was indicated by *** for p < 0.001, ** for p < 0.01, * for p < 0.05, and ns = not significant (p > 0.05).

## Results

### Generation of passage-able Exp-HLOs

iPSCs were differentiated into day 15 early hepatic progenitor cells using retinoic acid (RA) by following the Ouchi protocol^13^. Since the early hepatic progenitor cells express *LGR5* adult stem cell marker and *EPCAM* ductal epithelial marker, they were cultured in the expansion medium (EM) to form Exp-HLOs (**Figure 1**). The EM supports the expansion of bipotent ductal cells by cAMP activation (using Forskolin), TGF-β inhibition (A8301), and Wnt signaling (R-spondin1 or Rspo1). In the first attempt to generate Exp- HLOs from day 15 hepatic progenitor cells, we encountered several issues including slow organoid formation as well as the degradation of Exp-HLOs during passage and culture. Thus, we optimized the EM to promote the expansion of LGR5^+^ stem cells using Noggin^22,23^ and Wnt signaling throughout the expansion (**Supplementary Figure 2**). In addition, R-spondin1 purchased from R&D Systems (Rspo1_R&D) in the EM was replaced with home-made R-spondin1 conditioned medium (Rspo1_HM) to evaluate the efficacy of Rspo1_HM. Furthermore, based on precedent studies on critical growth factors necessary for HLO passage, we cultured Exp-HLOs in the EM without R-spo1 and EGF (EM – Rspo1 – EGF). As anticipated, Exp-HLOs were successfully generated in the EM containing either Rspo1_HM or Rspo1_R&D. It has been known that EGF mediates the ductal cell formation and promotes cell proliferation^24,25^. Interestingly, the EM – Rspo1 – EGF medium condition was also successful for robust organoid passage once Exp-HLOs had formed. Thus, we compared the gene expression level of the *LGR5* adult stem cell marker and *AFP* hepatic progenitor marker in Exp-HLOs cultured in the EM with Rspo1 and without Rspo1 to better understand the role of Rspo1 (**Supplementary Figure 3**). Since Rspo1 is known to support the proliferation of adult stem cells by activating Wnt/β-catenin signaling^26^, Exp-HLOs cultured in the EM with Rspo1 showed higher level of *LGR5* and *AFP* expression. With this result, the passage of Exp-HLOs were performed by mechanical dissociation using a pipette (**Supplementary Figure 4**) and enzymatic dissociation using TrypLE™ enzyme (**Figure 1C**). Since mechanical dissociation resulted in more variation in size and comparatively lower efficiency in the generation of Exp-HLOs (data not shown), we decided to use enzymatic dissociation for passage. The Exp-HLOs were successfully passaged up to 10 times by enzymatic dissociation and demonstrated similar morphology throughout the passages (**Figure 1C**). This result does not imply that the Exp-HLOs cannot be passaged beyond 10 times; rather, characterization beyond passage 10 has not yet been performed. The Exp-HLOs showed bipotent ductal epithelial characteristics. The expression levels of *LGR5* adult stem cell marker, *EPCAM* ductal epithelial marker, and *SOX9* ductal marker were maintained throughout the passages with few fluctuations, which were acceptable within experimental error ranges (**Figure 1D**). In addition, the EpCAM^+^ cell population was consistent within the range of 93 - 98% at different passage numbers (**Figure 1E**). The immunofluorescence staining of Exp-HLOs showed the expression of CK18 hepatocyte marker, EpCAM ductal epithelial marker, Ecad epithelial marker, along with no expression of ALB albumin, a functional biomarker of hepatocytes (**Figure 1F**). The uniformity of Exp-HLOs can be crucial to alleviate the issue of batch-to-batch variation in organoid generation, which was assessed both genotypically and phenotypically (**Figure 1**). Furthermore, the Exp-HLOs were cryopreserved and reseeded successfully to examine the feasibility of long-term storage (**Supplementary Figure 5A**). The morphology of Exp-HLOs did not change before and after cryopreservation. The P5 Exp-HLOs, thawed after 4 months of cryopreservation, proliferated after a 6-day lag phase for stabilization, as measured by the Cell Counting Kit 8 (**Supplementary Figure 5B**).

**Figure 1.**
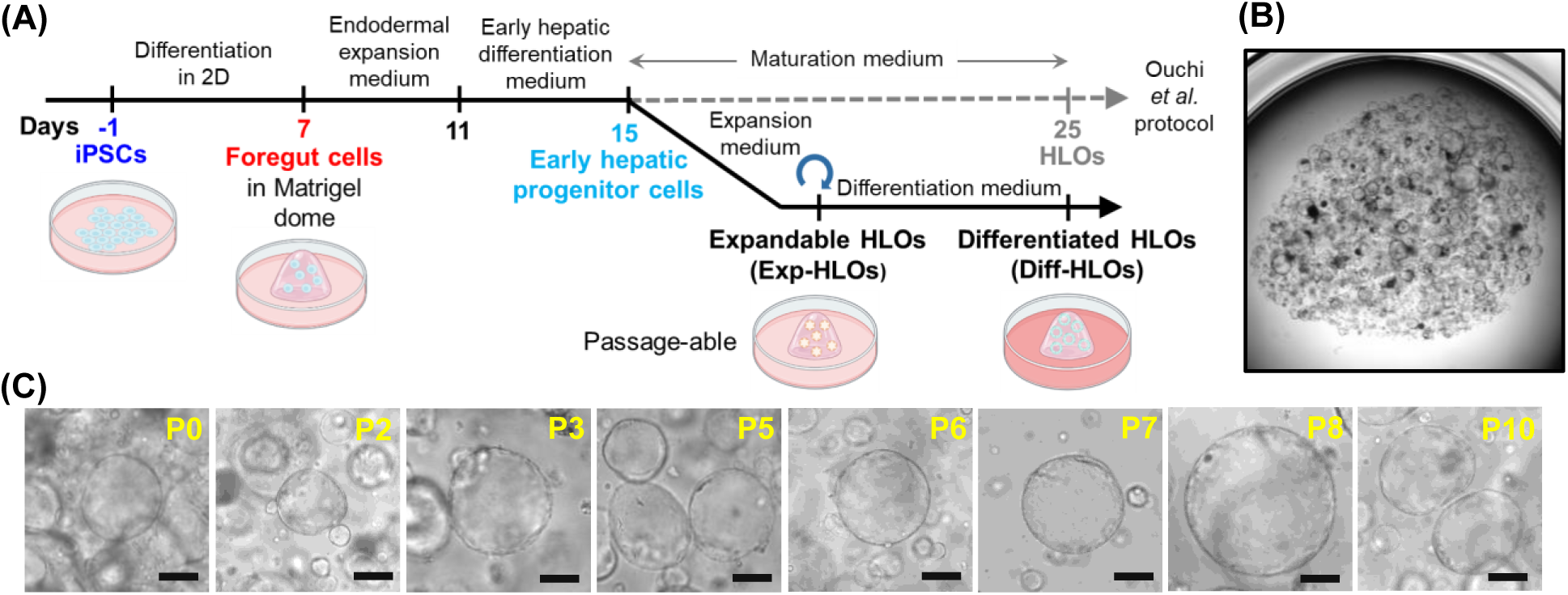
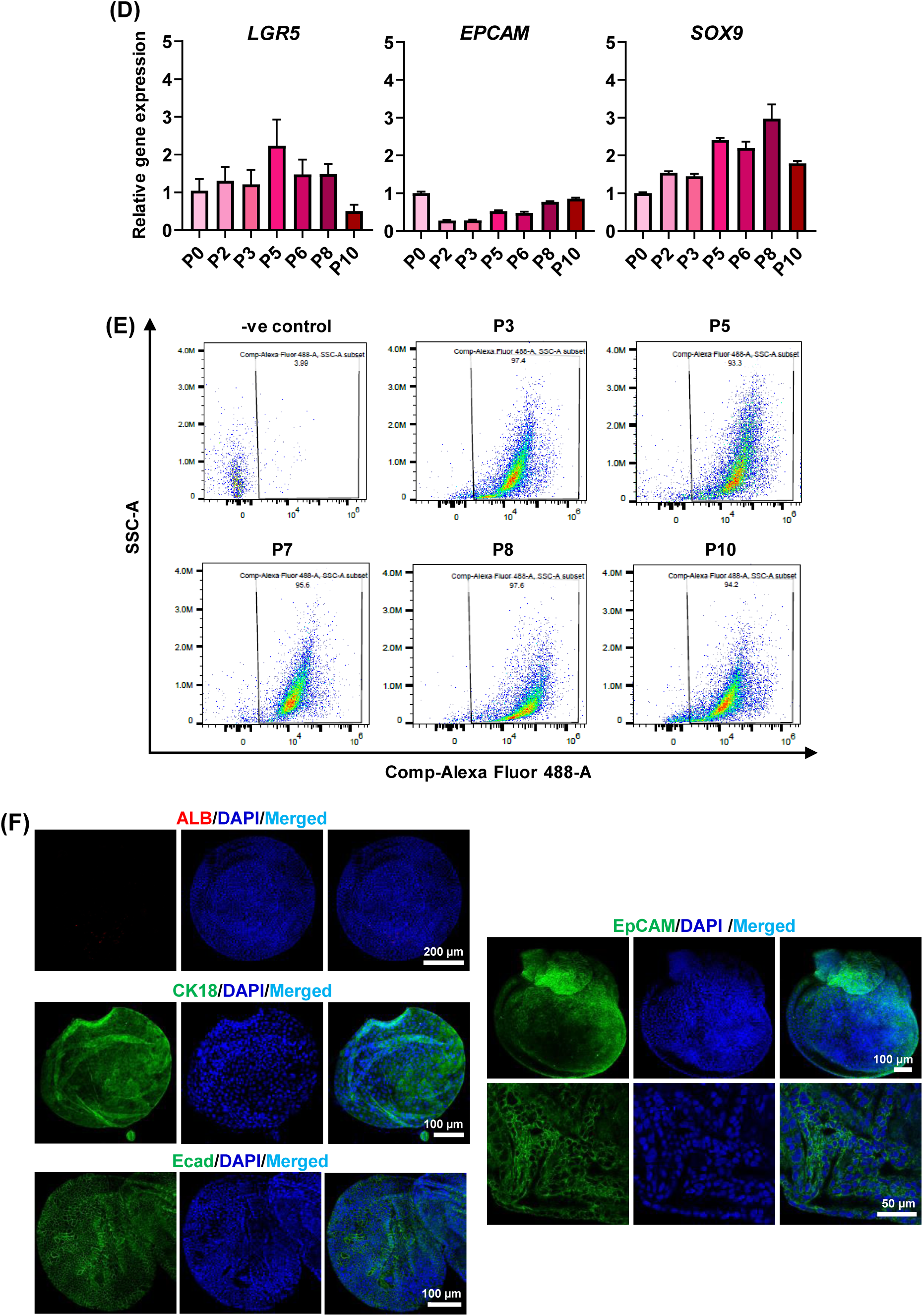
Generation of expandable human liver organoids (Exp-HLOs): **(A)** The differentiation protocol of iPSCs into Exp-HLOs and Diff-HLOs. **(B)** Bright-field image of a whole Matrigel dome containing Exp-HLOs in a 24-well plate. **(C)** Morphology of Exp-HLOs at different passage numbers. Scale bars: 200 µm. **(D)** Changes in hepatic gene expression in Exp-HLOs at different passage numbers, including *LGR5* adult stem cell marker, *EPCAM* ductal epithelial marker, and *SOX9* cholangiocyte marker. n = 4. One-way ANOVA was performed for statistical analysis. **(E)** The percentage of subpopulation expressing EpCAM ductal epithelial marker in Exp-HLOs at different passage numbers obtained by flow cytometry analysis. **(F)** Immunofluorescence staining of Exp-HLOs showing the expression of ALB albumin marker, CK18 hepatocyte marker, Ecad epithelial marker, and EpCAM ductal epithelial marker. Scale bars: 50 µm, 100 µm, and 200 µm.

### Differentiation of Exp-HLOs into Diff-HLOs

The Exp-HLOs were differentiated into Diff-HLOs using a differentiation medium (DM) containing the notch inhibitor DAPT, HGF, BMP7, and dexamethasone (**Figure 2A**). The Diff-HLOs were smaller and denser than Exp-HLOs (**Figures 2B and 2C**). The expression levels of stem cell and hepatic genes, including *LGR5* adult stem cell marker, *ALB* albumin marker, *ASGR1* hepatocyte marker, *EPCAM* ductal epithelial marker, and *AFP* hepatic progenitor marker, in (I) iPSCs, (II) day 15 early hepatic progenitor cells, (III) Exp-HLOs at P0, and (IV) day 10 Diff-HLOs at P1 were measured by qPCR analysis (**Figure 2D**). The expression of the *AFP* marker in iPSCs was not included due to its extremely low expression. The expression levels of *LGR5* and *EPCAM* genes were high in early hepatic progenitor cells and Exp-HLOs but decreased in Diff-HLOs. On the other hand, the expression levels of *ALB*, *ASGR1*, and *AFP* genes were higher in Diff-HLOs compared to Exp-HLOs, indicating the progression towards matured hepatic state after differentiation. However, the lower expression of mature hepatocyte marker *ASGR1* and higher expression of immature hepatocytes *AFP*, compared to Day 15 early hepatic progenitor cells indicate that 10 days of differentiation was insufficient for HLO maturation. The immunofluorescence staining of day 10 Diff- HLOs showed the expression of ALB albumin marker, Ecad epithelial marker, and CK18 hepatocyte marker (**Figure 2E**). Additionally, the Diff-HLOs demonstrated the capability of accumulating the fluorescently labeled bile acid in the canaliculus-like lumen structure at the center, indicating the function of BESP (**Figure 2F**). Furthermore, the Diff-HLOs showed the higher activity of CYP3A4 compared to Exp-HLOs (**Figure 2G**). Finally, the expression levels of hepatic genes, including *ALB* albumin marker, *ASGR1* hepatocyte marker, *SOX9* cholangiocyte marker, *CD68* Kupffer cell marker, *AFP* hepatic progenitor marker, and drug metabolizing enzymes such as *CYP3A4*, *UGT1A1,* and *SULT2A1*, in (I) day 25 HLOs obtained by using Ouchi et al. protocol^13^ and (II) day 10 Diff-HLOs were compared (**Figure 2H**). Interestingly, the HLOs generated by the two protocols were comparable, with a marginal increase in the expression level of drug metabolizing enzymes such as *CYP3A4* and *UGT1A1* from our protocol. It is worth noting that in the present work, all qPCR analysis has been performed on HLOs in conjunction with the mesenchyme population, whereas Ouchi et al.^13^ harvested and isolated HLOs to assess organoid maturity.

**Figure 2.**
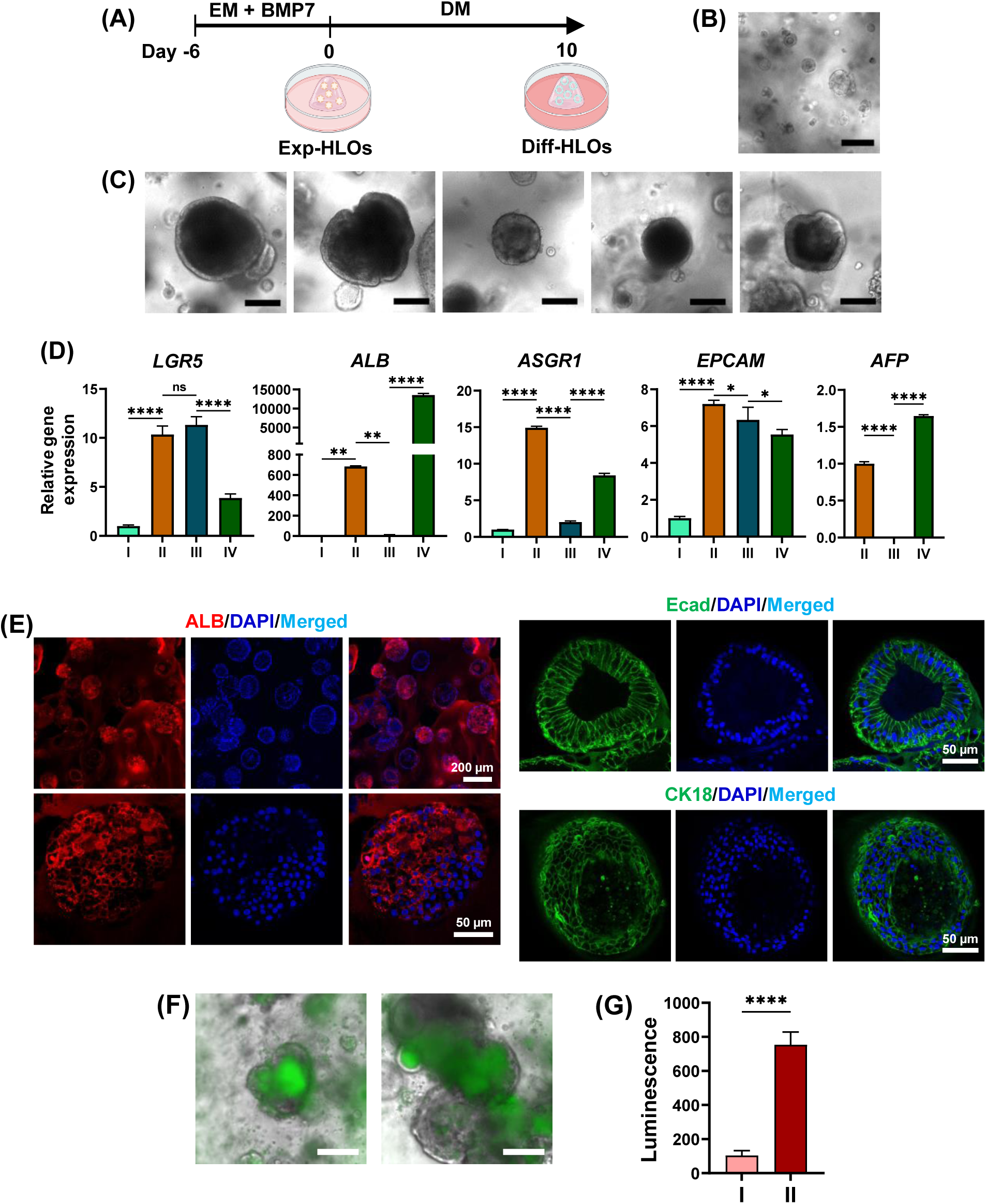
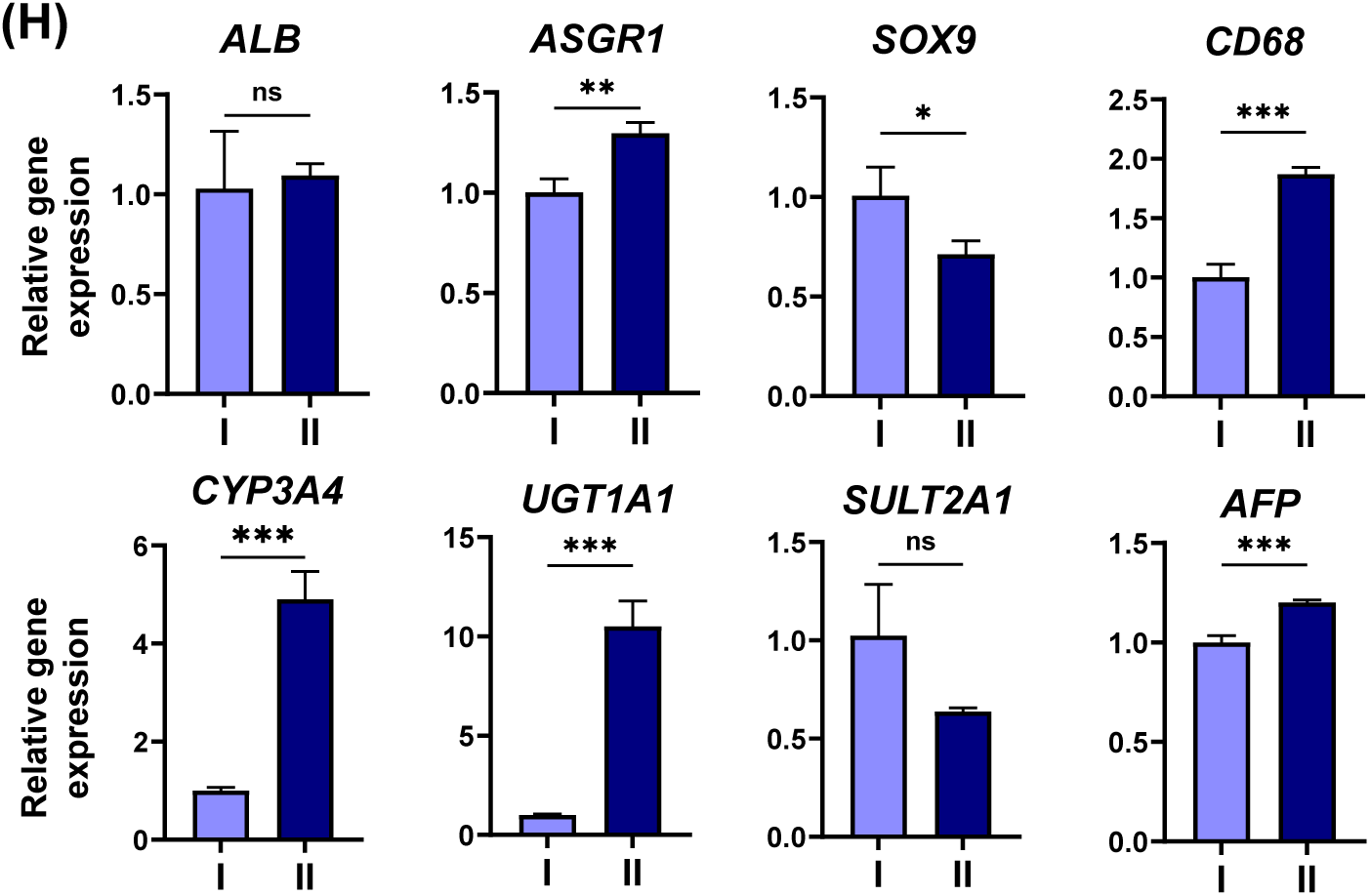
Generation of differentiated human liver organoids (Diff-HLOs): **(A)** The differentiation protocol of Exp-HLOs into Diff-HLOs by using the DM for 10 days. **(B)** Bright-field image of day 10 Diff- HLOs in Matrigel dome in a 24-well plate. Scale bar: 500 µm. **(C)** Representative morphology of day 10 Diff-HLOs. Scale bars: 200 µm. **(D)** Changes in hepatic gene expression in **(I)** iPSCs, **(II)** day 15 early hepatic progenitor cells, **(III)** Exp-HLOs at P0 and **(IV)** day 10 Diff-HLOs at P1, including *LGR5* adult stem cell marker, *ALB* albumin marker, *ASGR1* hepatocyte marker, *EPCAM* ductal epithelial marker, and *AFP* hepatic progenitor marker. n = 5. The significance of gene expression differences among the samples was analyzed using one-way ANOVA. **(E)** Immunofluorescence staining of day 10 Diff-HLOs showing the expression of ALB albumin marker, Ecad epithelial marker, and CK18 hepatocyte marker. Scale bars: 50 µm and 200 µm. **(F)** Uptake of fluorescein-labeled bile acid, cholyl-lysyl-fluorescein (CLF). Scale bars: 200 µm. **(G)** CYP3A4 activity in **(I)** Exp-HLOs and **(II)** day 10 Diff-HLOs measured by P450-Glo™ CYP3A4 luminescence assay kit. n = 9. The significance of CYP3A4 activity difference was analyzed by Student’s t-test. **(H)** Comparison of hepatic gene expression in **(I)** day 25 HLOs obtained by using Ouchi et al. protocol and **(II)** day 10 Diff-HLOs, including *ALB* albumin marker, *ASGR1* hepatocyte marker, *SOX9* cholangiocyte marker, *CD68* Kupffer cell marker, *AFP* hepatic progenitor marker, and drug-metabolizing enzymes such as *CYP3A4*, *UGT1A1,* and *SULT2A1*. n = 4. The significance of gene expression difference was analyzed by Student’s t-test. Both HLOs were generated in 50 µL Matrigel domes. Exp-HLOs at P3 were used to generate day 10 Diff-HLOs.

To further improve the maturity of Diff-HLOs, the differentiation medium (DM) was optimized in the presence and absence of 50 ng/ml EGF and 20 ng/mL OSM (**Figure 3A**). Interestingly, the DM without EGF and with OSM (DM – EGF + OSM) further enhanced the maturity of Diff-HLOs as indicated by the considerably enhanced expression of hepatic biomarkers including *ALB* albumin and *ASGR1* hepatocytes markers (**Figure 3B**). It is known that OSM induces hepatic maturation^27,28^, and thus has been supplemented in HLO maturation media previously. Similarly, EGF/EGF receptor (EGFR) is known to induce Notch1 that suppress hepatocyte commitment^25^.

**Figure 3.**
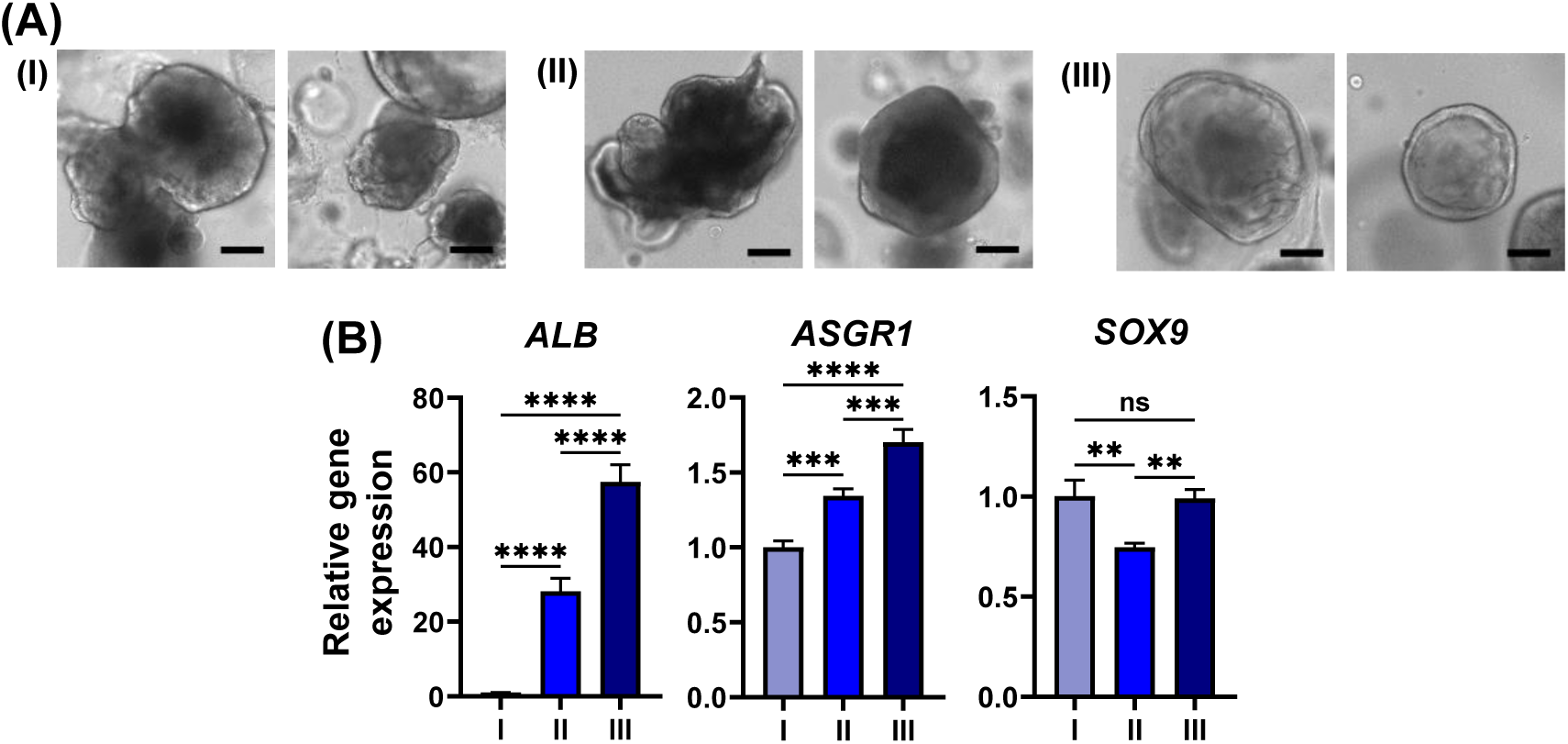
Optimization of the differentiation medium (DM) of Diff-HLOs. **(A)** Representative images of day 10 Diff-HLOs cultured in **(I)** the DM, **(II)** the DM without EGF, and **(III)** the DM without EGF and with OSM. Scale bars: 200 μm. **(B)** Comparison of hepatic gene expression in Diff-HLOs cultured in the three DM conditions, including *ALB* albumin marker, *ASGR1* hepatocyte marker, and *SOX9* cholangiocyte marker. n = 4. The significance of gene expression differences among the samples was analyzed using one-way ANOVA. Exp-HLOs at P4 were used to generate day 10 Diff-HLOs.

### Dynamic culture of Diff-HLOs in the pillar/perfusion plate

To further enhance the maturity of HLOs, we adopted an engineering approach and dynamically cultured Diff-HLOs in the pillar/perfusion plate (**Figures 4A and 4B**). We hypothesized that organoid maturity could be enhanced when cultured in a fluidic system due to higher diffusion of nutrients and oxygen into the core of organoids and fluidic shear stress. For assay miniaturization and high-throughput organoid culture, the 144PillarPlate was introduced to load Exp-HLOs in Matrigel on a 12 x 12 array of pillars (**Figure 4A**). Additionally, the complementary 144PerfusionPlate was introduced to generate bi-directional flow of the differentiation medium under the pillars and culture organoids in a dynamic condition (**Figure 4B**). By sandwiching the 144PillarPlate with Exp-HLOs onto the 144PerfusionPlate, Diff-HLOs can be generated in 12 different fluidic channels, each consisting of one upper reservoir (UR), one lower reservoir (LR), and a row of 12 perfusion wells connected by microchannels (**Figure 4C**). Gravity-driven fluid flow in forward and backward directions was generated in the 144PerfusionPlate coupled with the 144PillarPlate on a digital rocker (**Supplementary Figure 6**), which was simulated with SolidWorks at 1300 μL cell culture medium per fluidic channel, 10° tilting angle, and 1 minute of tilting angle change (**Figure 4D**). The maximum volumetric flow rate calculated was 10 μL/seconds, while the negative flow rates indicate the backward direction of flow. The volumetric flow rates in the pillar/perfusion plate are 100 – 1,000-fold faster than those obtained from conventional microfluidic devices and better mimic blood flow rates in large capillaries^20^. Additionally, the average shear stress on the surface of the pillars due to di-directional fluid flow simulated by SolidWorks was in the range of 0.2 - 0.5 dyne/cm^2^ (**Figure 4E**). The flow rate and shear stress can be controlled mainly by changing the tilting angle, as demonstrated in our previous study^20^. Nonetheless, the optimization of the flow rate and shear stress has not been performed in the present work to further improve the maturity of HLOs.

**Figure 4.**
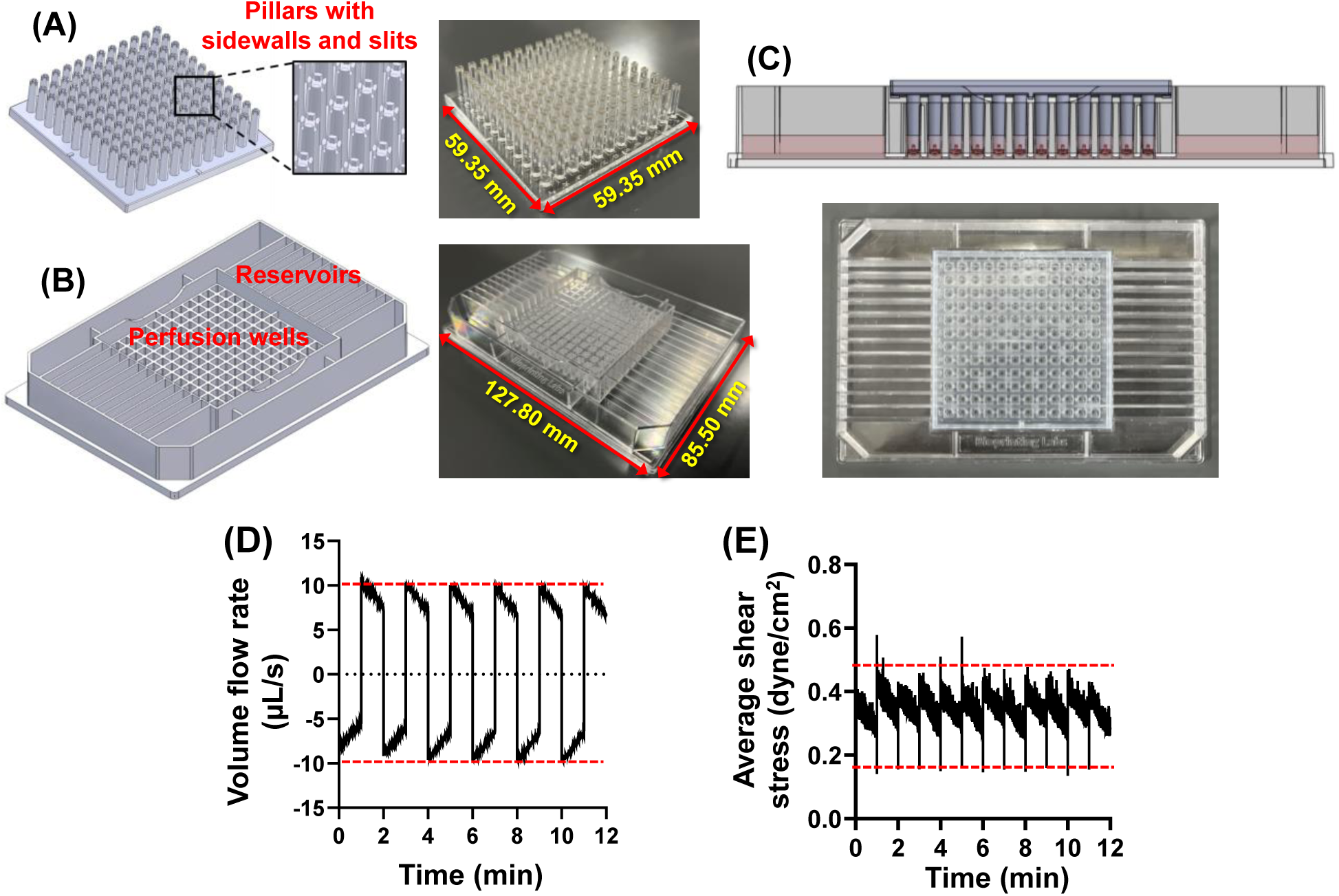
The injection-molded pillar plate and the complementary perfusion plate for dynamic organoid culture. **(A)** The 144PillarPlate with a 12 x 12 array of pillars for loading cells in hydrogels. **(B)** The 144PerfusionPlate with a 12 x 12 array of perfusion wells and a 2 x 12 array of reservoirs for supplying cell culture media. **(C)** The 144PillarPlate sandwiched onto the 144PerfusionPlate for dynamic cell culture. The 144PerfusionPlate contains 12 fluidic channels, each channel consisting of one upper reservoir (UR), one lower reservoir (LR), and a row of 12 perfusion wells connected by microchannels. **(D)** Volumetric flow rates in forward and backward directions in the 144PerfusionPlate coupled with the 144PillarPlate simulated with SolidWorks at 1300 μL cell culture medium per fluidic channel, 10° tilting angle, and 1 minute frequency of tilting angle change. The negative flow rates indicate the opposite direction of flow. **(E)** Average shear stress on the surface of the pillars by fluid flow simulated by SolidWorks.

To demonstrate dynamic culture of Diff-HLOs in the pillar/perfusion plate, Exp-HLOs generated in the Matrigel dome in the 24-well plate were enzymatically dissociated, isolated by centrifugation, resuspended in fresh Matrigel, and then printed on the 144PillarPlate by using a 3D bioprinter. The enzymatically dissociated cell clumps suspended in Matrigel were uniformly printed on the 144PillarPlate with a CV value of 10% (**Supplementary Figure 7A**). The 144PillarPlate with cell clumps was sandwiched onto the 144PerfusionPlate containing the EM supplemented with BMP7 and cultured for 6 days to form Exp-HLOs (**Supplementary Figure 7B**). The Exp-HLOs on the pillar plate were further cultured in the DM supplemented with OSM for up to 20 days in a dynamic condition in the perfusion plate (**Supplementary Figure 7C**). The Diff-HLOs developed a denser and larger morphology over time, and the expression levels of critical hepatic biomarkers, including *ALB*, *ASGR1*, *SOX9*, *CD68*, *CYP3A4*, *UGT1A1*, and *SULT2A1*, were significantly enhanced with a long duration of differentiation (**Supplementary Figure 7D**). This result represents a significant advancement, as iPSC-derived HLOs generated by the Ouchi et al. protocol^13^ typically exhibited gradual loss of hepatic functions post-maturation (**Supplementary Figure 8**).

Compared to day 20 Diff-HLOs statically cultured in the 144PillarPlate/384DeepWellPlate, day 20 Diff-HLOs dynamically cultured in the 144PillarPlate/144PerfusionPlate exhibited denser cell morphology with almost double in ATP content (**Figures 5A, 5B, and 5F**). Moreover, the expression levels of critical hepatic biomarkers, including *ALB*, *ASGR1*, *SOX9*, *VM*, *CD68*, *CYP3A4*, *UGT1A1*, and *SULT2A1*, were significantly higher in dynamically cultured Diff-HLOs compared to their static counterparts, highlighting the importance of high-throughput, dynamic organoid culture (**Figure 5C**). The immunofluorescence staining of day 20 Diff-HLOs showed the expression of ALB albumin marker comparatively enhanced in the dynamic culture condition compared to the static culture condition (**Figures 5D and 5E**). Besides the improvements in maturity and functionality, the generation of Diff-HLOs in dynamic culture also showed uniformity, with CV value of 21.5% based on ATP-based viability assessment (**Figure 5F**).

**Figure 5.**
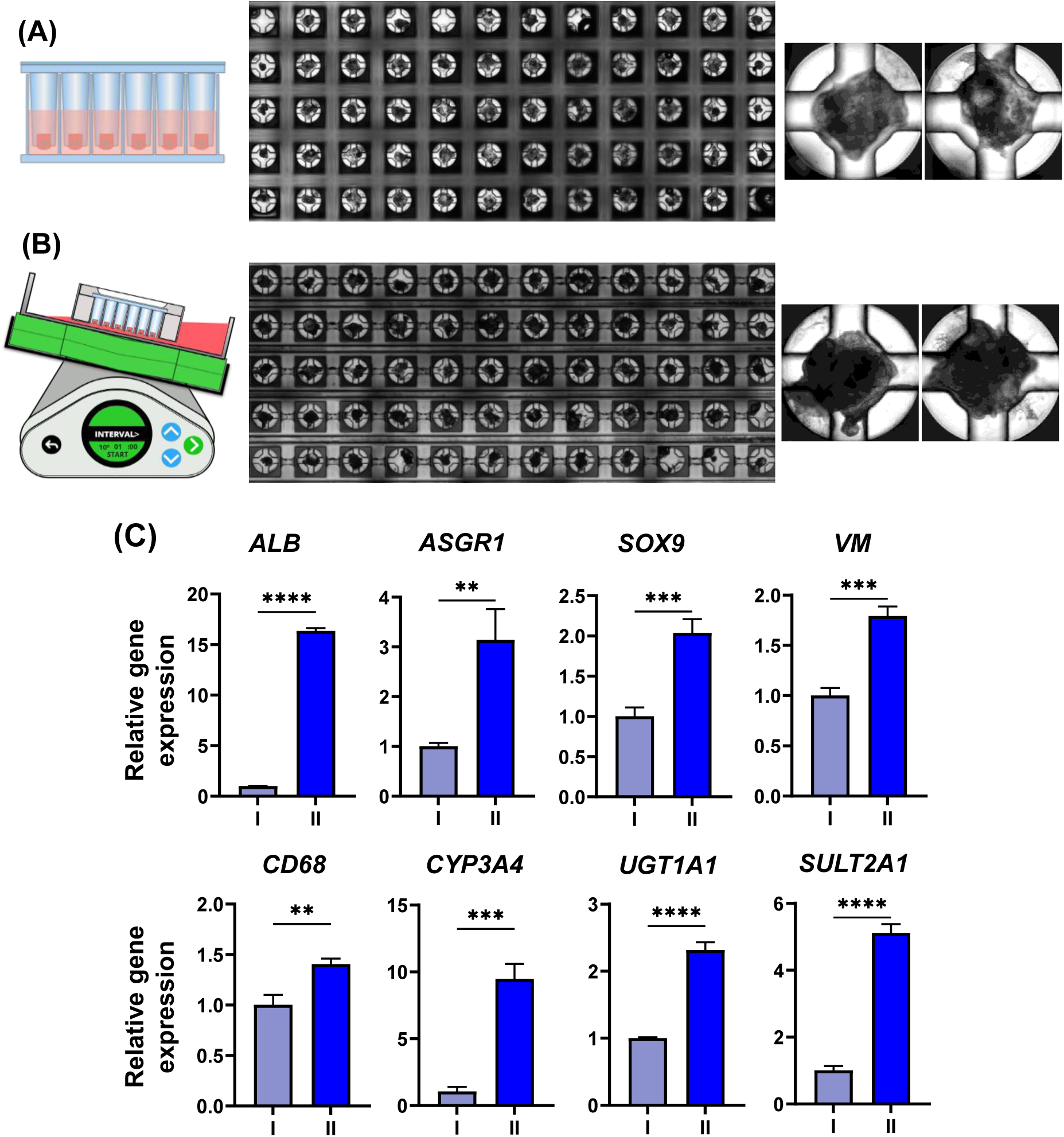
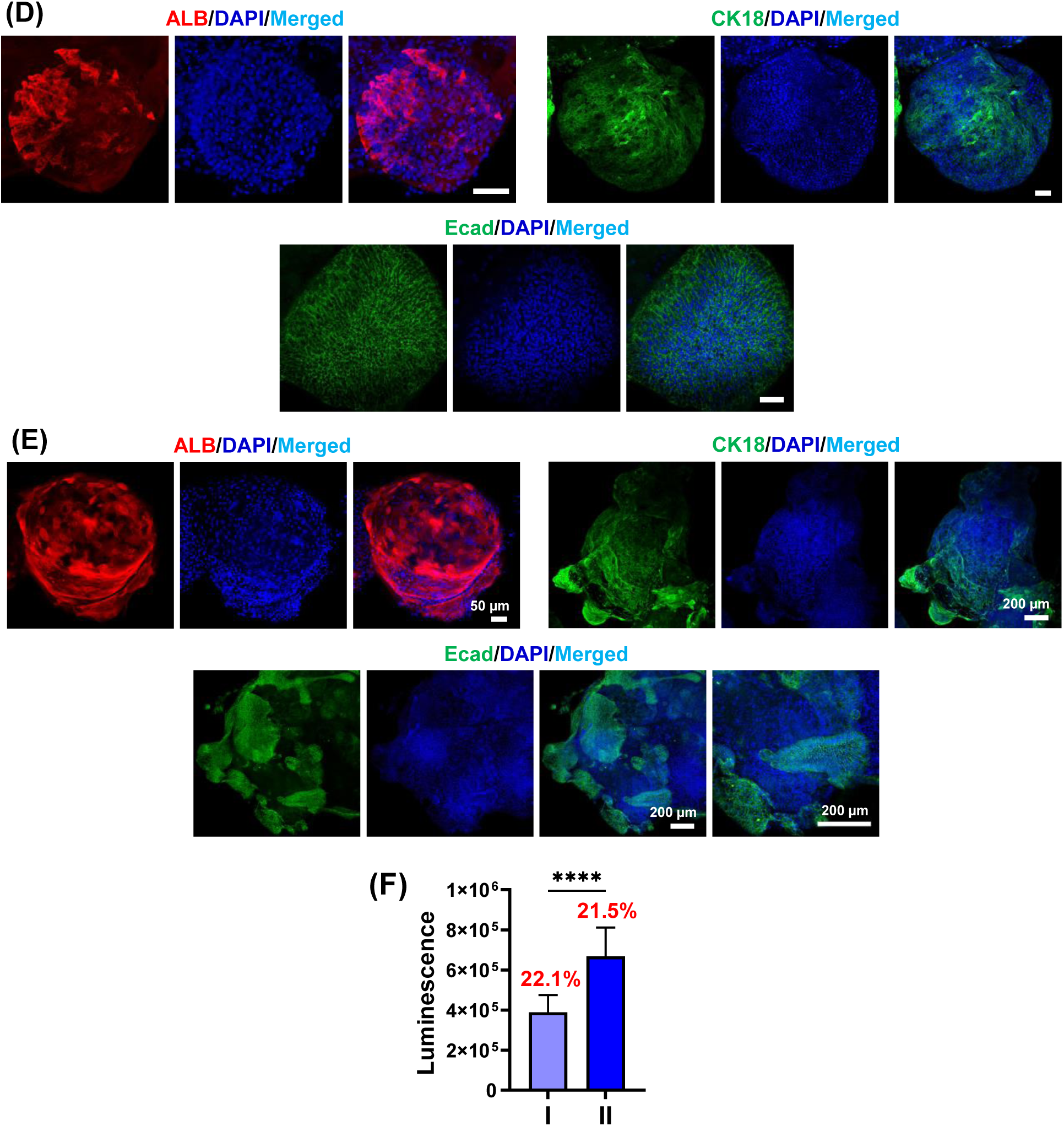
Static and dynamic culture of Diff-HLOs on the pillar plate. **(A)** Stitched (left) and representative (right) images of day 20 Diff-HLOs cultured in a static condition using the 144PillarPlate coupled with the 384DeepWellPlate. **(B)** Stitched (left) and representative (right) images of day 20 Diff- HLOs cultured in a dynamic condition using the 144PillarPlate coupled with the 144PerfusionPlate. **(C)** Comparison of hepatic gene expression in day 20 Diff-HLOs cultured in **(I)** static and **(II)** dynamic conditions, including *ALB*, *ASGR1*, *SOX9*, *VM*, *CD68*, *CYP3A4*, *UGT1A1*, and *SULT2A1*. n = 72. The significance of gene expression difference was analyzed by Student’s t-test. **(D)** Immunofluorescence staining of day 20 Diff-HLOs cultured in a static condition. Scale bars: 50 µm. **(E)** Immunofluorescence staining of day 20 Diff-HLOs cultured in a dynamic condition. Scale bars: 50 µm (multiple HLOs) and 200 µm (single HLOs on a pillar). **(F)** Uniform generation of day 20 Diff-HLOs on both **(I)** static and **(II)** dynamic culture conditions analyzed using ATP-based cell viability assay kit with CV values of 22.1% and 21.5%, respectively. n = 60 for both culture conditions. Exp-HLOs at P5 were used to generate day 20 Diff- HLOs.

### Assessment of drug-induced liver injury (DILI) by using Diff-HLOs in the pillar/perfusion plate and the recovery of the liver from DILI

The 144PillarPlate with 144 replicates of Diff-HLOs was sandwiched onto the 144PerfusionPlate containing six different concentrations of a compound to obtain a dose-response curve with 24 replicates of Diff-HLOs per concentration (**Figure 6A**). Acetaminophen and troglitazone were used as model compounds for drug-induced liver injury (DILI) within the concentration range of 0 – 10,000 μM for acetaminophen and 0 – 500 μM for troglitazone (**Figures 6B and 6C**). After 3 days of dynamic compound treatment, the IC_50_ value obtained for acetaminophen was 4,060 μM whereas the IC_50_ value of troglitazone was 164.6 μM. Acetaminophen is a widely used over-the-counter (OTC) analgesic and antipyretic medication for relieving mild-to-moderate pain and fever. However, it can cause acute liver injury and death at high doses^29^. From recent studies, the IC_50_ value of acetaminophen is reported to be 4,286 µM with gold-standard, primary human hepatocytes (PHHs) and 4,036 µM with hepatocyte alternative, HepaRG cells^30^, both of which are similar to the IC_50_ obtained from this study. In addition, troglitazone was approved for the treatment of type 2 diabetes, but already withdrawn from the market due to cases of acute liver failure^31^. Recent studies reported that the IC_50_ value for troglitazone is 57 µM with PHHs and 45 µM with HepaRG cells^30^, both of which are within the typical one dose error range. The predictivity of DILI with Diff-HLOs cannot be assessed accurately with the two model compounds. A set of DILI compounds with different mechanisms of action should be used to assess the potential of Diff-HLOs to replace PHHs or HepaRG cells.

**Figure 6.**
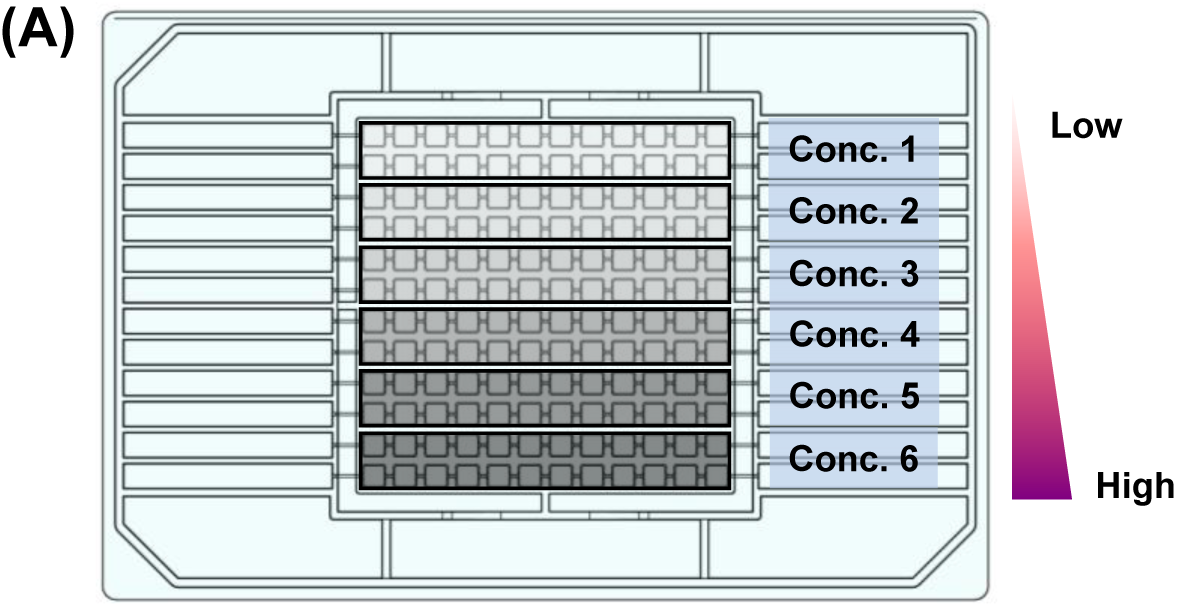
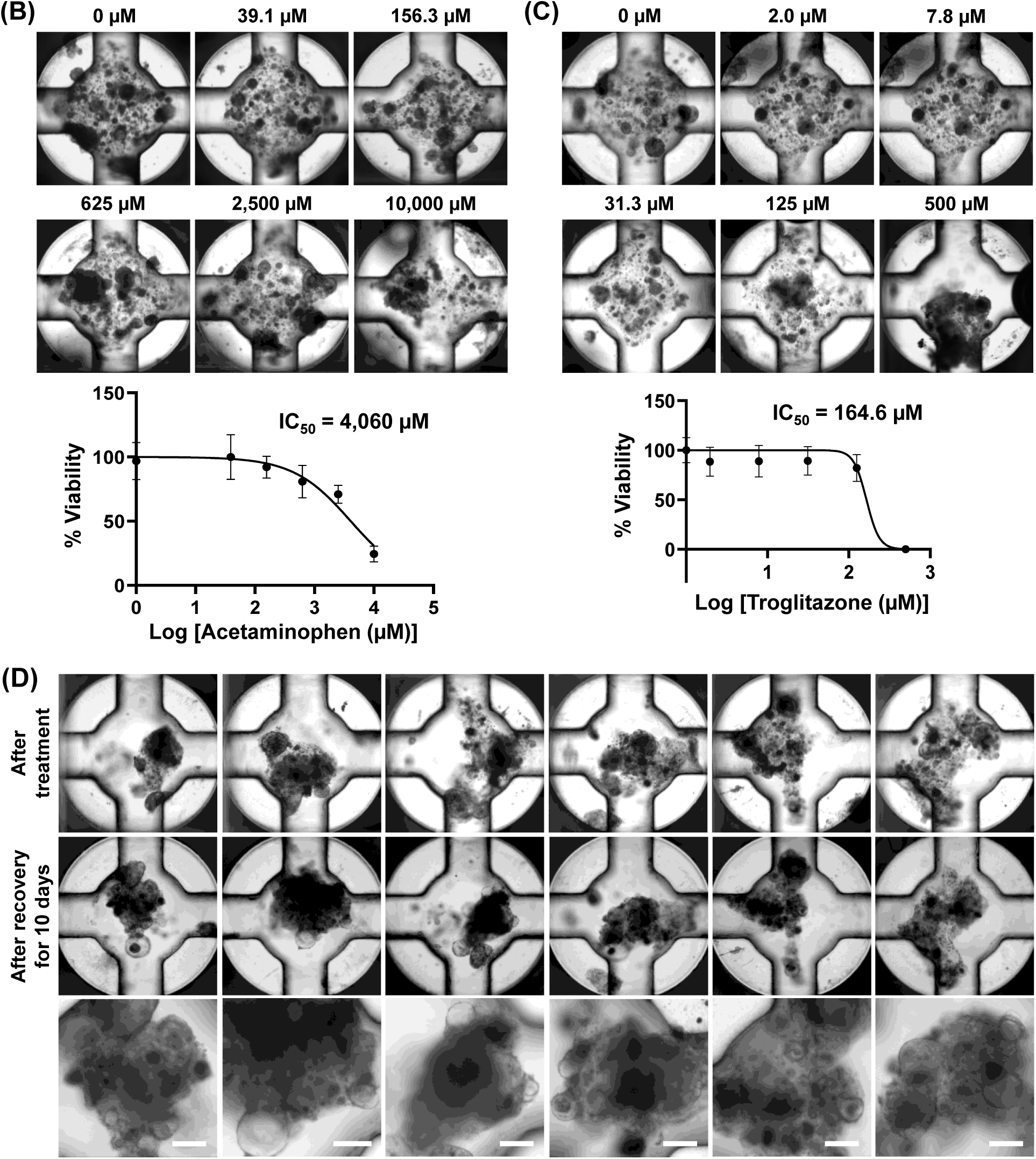
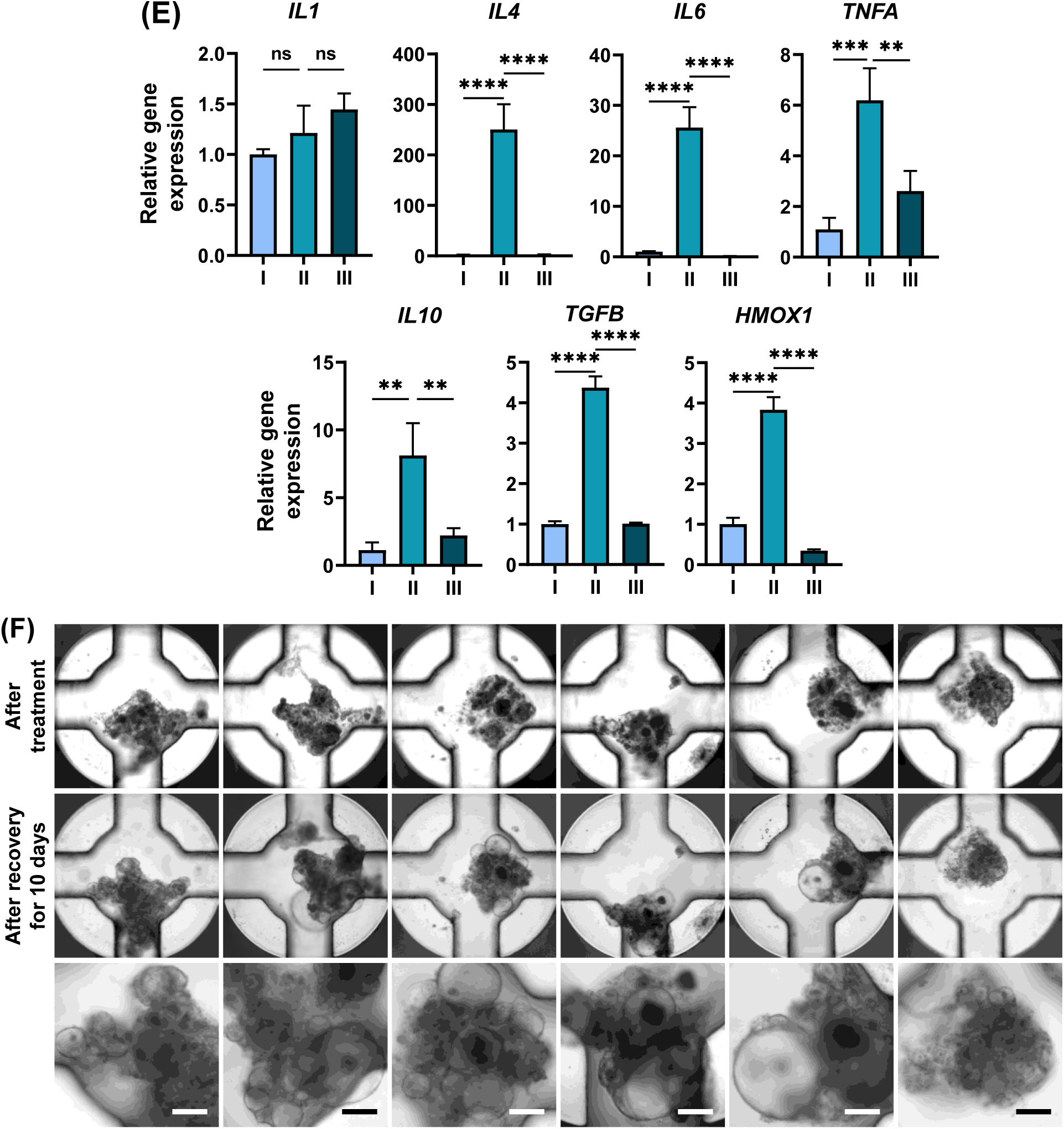
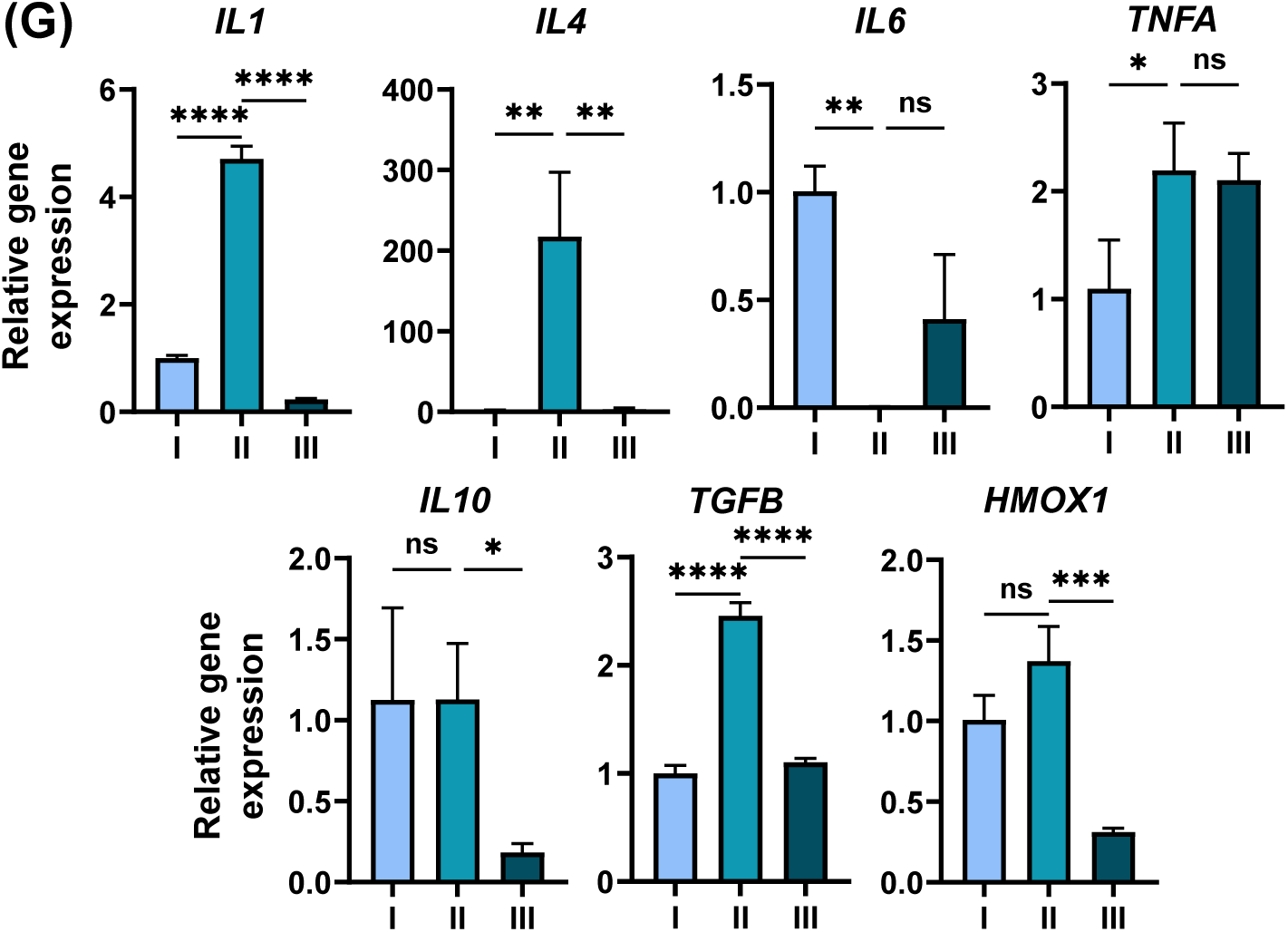
Assessment of hepatotoxicity with model compounds and liver recovery after compound treatment using day 20 Diff-HLOs in the pillar/perfusion plate: **(A)** Schematic of compound dosage in the 144PerfusionPlate. **(B)** Representative images of Diff-HLOs after acetaminophen treatment for 3 days at the concentration range of 0 – 10,000 μM and its dose response curve. **(C)** Representative images of Diff- HLOs after troglitazone treatment for 3 days at the concentration range of 0 – 500 μM and its dose response curve. For (B) and (C), Exp-HLOs at P7 were differentiated into Diff-HLOs. **(D)** Representative images of Diff-HLOs after 3 days of treatment with 5,000 μM acetaminophen (top panel) and after 10 days of recovery in the EM (middle and bottom panels). Scale bars: 200 μm. **(E)** Changes in inflammatory gene expression in **(I)** Diff-HLOs (control), **(II)** Diff-HLOs after 3 days of treatment with 5,000 μM acetaminophen, and **(III)** Diff-HLOs after 10 days of recovery in the EM, including *IL1*, *IL4*, *IL6, TNFA, IL10, TGFB,* and *HMOX1*. n = 12. The significance of gene expression difference among the samples was analyzed using one-way ANOVA. **(F)** Representative images of Diff-HLOs after 3 days of treatment with 125 μM troglitazone (top panel) and after 10 days of recovery in the EM (middle and bottom panels). Scale bars: 200 μm. **(G)** Changes in inflammatory gene expression in **(I)** Diff-HLOs (control), **(II)** Diff-HLOs after 3 days of treatment with 125 μM troglitazone, and **(III)** Diff-HLOs after 10 days of recovery in the EM, including *IL1*, *IL4*, *IL6, TNFA, IL10, TGFB,* and *HMOX1*. n = 12. The significance was analyzed by using one-way ANOVA. For (D) – (G), Exp-HLOs at P9 were differentiated into Diff-HLOs.

In addition to the assessment of DILI of model compounds, we evaluated liver recovery from DILI using Diff-HLOs in the pillar/perfusion plate, a process traditionally performed with animal models. Briefly, Diff-HLOs were dynamically treated for 3 days at their IC_50_ concentration (e.g., 5,000 μM for acetaminophen and 125 μM for troglitazone) to induce severe liver organoid damage (**Figures 6D and 6F**). This was followed by the dynamic culture of damaged Diff-HLOs in the EM for 10 days to regenerate HLOs. The EM induces the proliferation of hepatocytes and progenitor cells due to the presence of Rspo1, HGF, and Forskolin, which act as mitogens^32,33^, resulting in the regeneration of Diff-HLOs. After 10 days of recovery, we observed the expansion of Diff-HLOs, indicated by the appearance of circular organoids, which are the typical morphology of Exp-HLOs (**Figures 6D and 6F**). Furthermore, epithelial cells are known to produce pro-inflammatory cytokines to initiate the acute sterile inflammatory response, resulting from damage-associated molecular patterns (DAMPs) recognized by pattern-recognition receptors (PRRs)^33,34^. Interestingly, pro-inflammatory mediators such as *IL1*, *IL4, IL6,* and *TNFA* as well as *HMOX1* cellular stress marker were strongly expressed in Diff-HLOs after acetaminophen treatment (**Figure 6E**). Additionally, the expression levels of anti-inflammatory mediators such as *IL10* and *TGFB* were also enhanced, which are responsible for suppressing inflammation, inducing tissue remodeling, and facilitating tissue homeostasis^35^. A similar trend of pro- and anti-inflammatory marker expression, except for *IL6* and *IL10*, was observed for Diff-HLOs exposed to troglitazone (**Figure 6G**). Troglitazone, a thiazolidinedione drug, selectively binds to PPAR-γ and activates PPAR-γ, leading to the suppression of pro-inflammatory pathways, including IL6 production^36^. Since liver regeneration after DILI can be dose-dependent and significantly impaired beyond a certain dosage, the assessment of Diff-HLO recovery after compound treatment could provide valuable insights into *in vivo* liver damage and regeneration while estimating safe drug dosages for patient treatment^37^.

## Discussion

In 2015, Huch et al. generated functional liver organoids for the first time by the expansion and differentiation of LGR5^+^ adult bile duct-derived bipotent progenitor cells obtained from human liver biopsy samples^7^. Since then, several protocols have been published demonstrating the expansion and differentiation of functional liver organoids using either human embryonic stem cells (ESCs)^38,39^ or induced pluripotent stem cells (iPSCs)^38,40,41^. However, precedent studies mainly focused on hepatocyte functions, including albumin secretion and the expression of drug metabolizing enzymes, with only a few studies demonstrating the expression of cholangiocyte markers, indicating the presence of cholangiocyte-like cells in mature HLOs. None of the early studies focused on the cells of mesenchymal origin present in the liver, including Kupffer cells and stellate cells, along with endodermal hepatocytes and cholangiocytes necessary for modeling complex diseases such as fibrosis and alcoholic liver disease (ALD). One study performed the co-culture of expandable hepatic organoids with human fetal liver mesenchymal cells isolated from fetal liver tissues and demonstrated the improvement in hepatic functions, including albumin secretion, urea production, and CYP3A4 activity^39^. In 2019, Ouchi et al. demonstrated the generation of multicellular HLOs composed of hepatocyte-, cholangiocyte-, stellate-, and Kupffer-like cells by the differentiation of patient-derived iPSCs^13^. Although the Ouchi protocol is robust and allows the cryopreservation of foregut cells, it still requires a three-week-long multi-step differentiation and maturation process, which may lead to batch-to-batch variation. Additionally, mature HLOs gradually lose their hepatic functions over time (**Supplementary Figure 8**), which could be a critical issue for high-throughput screening (HTS) of hepatotoxic compounds. For the scale-up production of HLOs for compound screening, HLOs cultured in Matrigel domes need to be physically or enzymatically dissociated, which is labor-intensive and could lead to the loss of HLOs, damage to HLO structure and function, and batch-to-batch variation in their size. To resolve some of these issues and generate passage-able HLOs, we focused on day 15 early hepatic progenitor cells expressing *LGR5* from the Ouchi protocol^13^ and modified the differentiation protocol used in Huch et al.^7^ The day 15 early hepatic progenitor cells were successfully differentiated and expanded into Exp-HLOs that were genotypically and phenotypically stable throughout 10 passages (**Figure 1**). The Exp- HLOs were further differentiated into Diff-HLOs with enhanced hepatic functions, along with the presence of ductal features and the expression of cellular biomarkers of mesenchymal origin (**Figure 2**). Although Diff-HLOs obtained from our modified protocol showed comparatively higher expression levels of drug metabolizing enzymes such as *CYP3A4* and *UGT1A1* compared to those in HLOs from the Ouchi et al.^13^ protocol (**Figure 2H**), there remains potential for further enhancing HLO maturity and function. Among the few known compounds, the addition of OSM along with the removal of EGF in the DM improved the expression levels of critical hepatic genes, including *ALB* and *ASGR1* in Diff-HLOs (**Figure 3B**).

One of the engineering approaches for improving the maturity of organoids is the incorporation of micro-/milli-fluidic devices introducing fluid flow into the culture^42^. Several precedent studies have demonstrated that the shear stress due to fluid flow resulted in improved viability, vascularization, and maturity of the organoids^43–50^. The improvement in organoid maturity could be due to the significant role of biophysical cues such as shear stress in the organoid development^51,52^ and increased oxygenation and exposure to growth and differentiation factors in convective fluid flow transport^46,53^. Several companies have developed the state-of-the-art microfluidic organ-on-chip platforms, including Emulate’s Liver-Chip^54^, Javelin’s Liver Tissue Chip^55^, CN-Bio’s PhysioMimix^®^ Liver Plate^56^, Nortis’s ParVivo™ Microfluidic Chip^57^, to develop more predictive *in vitro* human liver models. However, these systems are inherently low throughout and expensive to operate due to the requirement for tubes, pumps, and proprietary incubator systems. In addition, some of these platforms are specifically designed to mimic a certain aspect of liver functions rather than culturing whole liver organoids.

Currently, organoid culture, compound testing, and organoid imaging and analysis have been performed on different platforms. For example, iPSCs were differentiated into mature HLOs in Matrigel domes in a 24-well plate. For high-throughput compound testing with HLOs, Matrigel domes were dissociated physically or enzymatically to harvest HLOs, leading to low throughput due to the manual steps involved. Harvested HLOs suspended in cell culture media were dispensed in a high-density microtiter well plate (e.g., 384-well plate) using a liquid handling system for cell-based assays^15^. This is one of key bottlenecks in organoid-based compound screening. Recently, Sun Bioscience developed the Gri3D^®^ microwell array plate^58^ for high-throughput organoid culture and imaging for compound testing. Although this is a significant advancement in the field, this platform only supports static culture of small organoids, which don’t require ECM encapsulation. To address these issues, we designed the pillar/perfusion plate platform and demonstrated rapid cell loading, organoid culture, compound testing, and organoid imaging and analysis in a single system^19,20,59–61^. In the present work, we developed the 144PillarPlate that allowed for the rapid and reproducible printing of 144 replicates of Exp-HLOs, which was coupled with the 144PerfusionPlate to generate 144 replicates of Diff-HLOs in 10 – 20 days of differentiation. Since HLOs were generated in a dynamic condition, there was no concern for the diffusion limitation of nutrients and oxygen, leading to high maturity of HLOs compared to their static counterparts (**Figure 5**). Additionally, unlike conventional iPSC-derived HLOs, Diff-HLOs showed higher maturity with longer culture time in the DM (**Supplementary Figure 7**). Compound testing with Diff-HLOs and organoid staining were performed in the pillar/perfusion plate in high throughput (**Figure 6**).

In addition to testing DILI potential of the model compounds, we demonstrated the recovery of the liver post-compound treatment using Diff-HLOs in the pillar/perfusion plate. Liver regeneration post-liver damage is crucial to better understanding the severity of DILI and can provide insights into patient survival after overdose and the repair mechanisms of the liver, potentially leading to new treatment options for patients with high vulnerability to DILI. In addition, the mechanisms of liver repair and regeneration post- DILI could be different from those after partial hepatectomy, which has been the main focus of previous studies on liver regeneration^32^. Traditionally, animal models such as rats and mice have been used to study dose-dependent, liver regeneration after drug treatment^62–64^. Recently, one study has demonstrated the capability of hepatocyte-like liver organoids to recapitulate liver regeneration after acetaminophen treatment^38^. Nonetheless, the liver organoids used were at an immature expandable stage, making them less sensitive to metabolism-sensitive compounds such as acetaminophen.

One of the most significant issues, often not discussed in detail, is the immaturity of HLOs. Significant improvements in HLO maturity and functionality have been made in the present work by developing Exp- HLOs, optimizing additives in the DM, employing long-term differentiation, and introducing dynamic organoid culture. The expression levels of key liver markers in dynamically cultured day 20 Diff-HLOs are higher than those in day 25 HLOs from the Ouchi et al.^13,15^ protocol (**Supplementary Figure 9**), even though the former includes the mesenchymal cell population. We achieved improved maturity and functionality compared to other previously published ASC- and iPSC-derived liver organoid protocols^7,38,40,65^. However, the maturity of Diff-HLOs still does not match that of *in vivo* liver tissue (**Supplementary Figure 9**). Overall, our dynamically cultured day 20 diff-HLOs in the pillar/perfusion plate exhibited a 10 – 100-fold lower maturity level compared to human adult liver tissues. Considering it takes approximately two years after birth for the human liver to attain full maturity^66^, it may be overly ambitious to expect HLOs to reach a maturity level directly comparable to liver tissue in just a few weeks of culture. A strategy involving the introduction of a cocktail of inducers for drug metabolizing enzymes during the maturation process could significantly enhance their expression to levels comparable with those of liver tissue. Furthermore, overexpressing transcription factors related to organoid maturity through gene regulatory network analysis^67^ could lead to a shorter organoid maturation period. Nonetheless, more efforts should be devoted to enhancing the maturity of HLOs to fully realize their potential in translational applications.

## Conclusions

In this study, we successfully generated passage-able and cryopreserve-able Exp-HLOs with genotypic and phenotypic stability throughout 10 passages, which were differentiated into functional Diff-HLOs in 10 - 20 days of culture. Additionally, we introduced the pillar plate to print and load 144 replicates of Exp-HLOs per plate, which was coupled with the perfusion plate to generate 144 replicates of Diff-HLOs in a dynamic condition. Our 3D bioprinting approach of passage-able liver organoids in the pillar/perfusion plate enabled high-throughput organoid culture and compound testing *in situ* in a dynamic condition, addressing several technical challenges in the organoid field. These challenges include user unfriendliness in dynamic organoid culture, high batch-to-batch variability, low throughput in organoid imaging, low organoid maturity, and high cost in organoid-based assays. The regenerative Diff-HLOs allowed us to test drug-induced liver injury (DILI) with model compounds as well as the recovery of the liver post-DILI, which has traditionally been tested with animal models. Thus, we envision that human organoids in the pillar/perfusion plate could be used for translational applications, including high-throughput compound screening, disease modeling, and precision medicine with cells from patients.

## Acknowledgements

This study was financially supported by the National Institutes of Health (NIEHS R43ES035653, NIDDK UH3DK119982, and NCATS R44TR003491) and the Ohio Third Frontier Commission (TVSF Phase IB and Phase II).

## Data availability statement

The data supporting this article has been included as part of the Supplementary Information.

## Supplementary information

**Supplementary Table 1.**
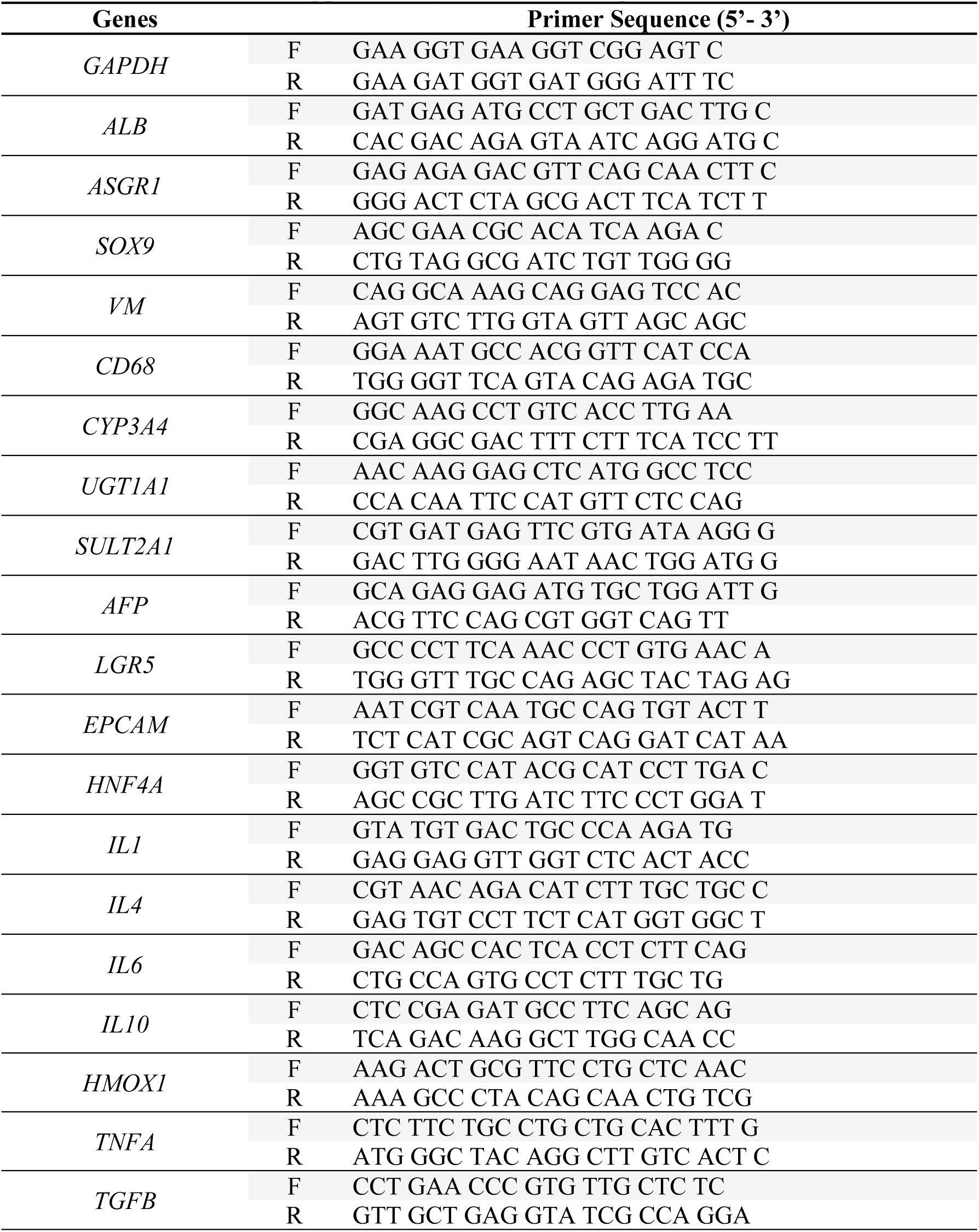
List of primers.

**Supplementary Table 2.** List of primary antibodies.

| Primary antibodies | Host/Isotype | Vendor | Dilution/Working concentration |
| --- | --- | --- | --- |
| ALB | Rabbit/polyclonal (IgG) | ab2406;<br>Abcam | 5 µg/mL |
| E-cad | Mouse/monoclonal (IgG<sub>1</sub>) | sc-21791; SantaCruz | 1:40 |
| VM | Rabbit/polyclonal (IgG) | SAB1305445;<br>Sigma-Aldrich | 1:40 |
| CK18 | Sheep/polyclonal (IgG) | AF7619;<br>R&D Systems | 5 µg/mL |
| EpCAM | Mouse/monoclonal (IgG<sub>1</sub>) | sc-25308; SantaCruz | 1:40 |

**Supplementary Table 3.** List of secondary antibodies.

| Secondary antibodies | Host | Vendor | Dilution |
| --- | --- | --- | --- |
| Donkey anti-Mouse IgG (H+L) Highly Cross-Adsorbed Secondary Antibody, Alexa Fluor™ 488 | Donkey | A-21202;<br>Invitrogen | 1:400 |
| Donkey anti-Rabbit IgG (H+L) Highly Cross-Adsorbed Secondary Antibody, Alexa Fluor 647 | Donkey | A-31573;<br>Invitrogen | 1:400 |
| Donkey anti-Sheep IgG (H+L) Highly Cross-Adsorbed Secondary Antibody, Alexa Fluor 488 | Donkey | A-11015;<br>Invitrogen | 1:400 |

**Supplementary Figure 1.**
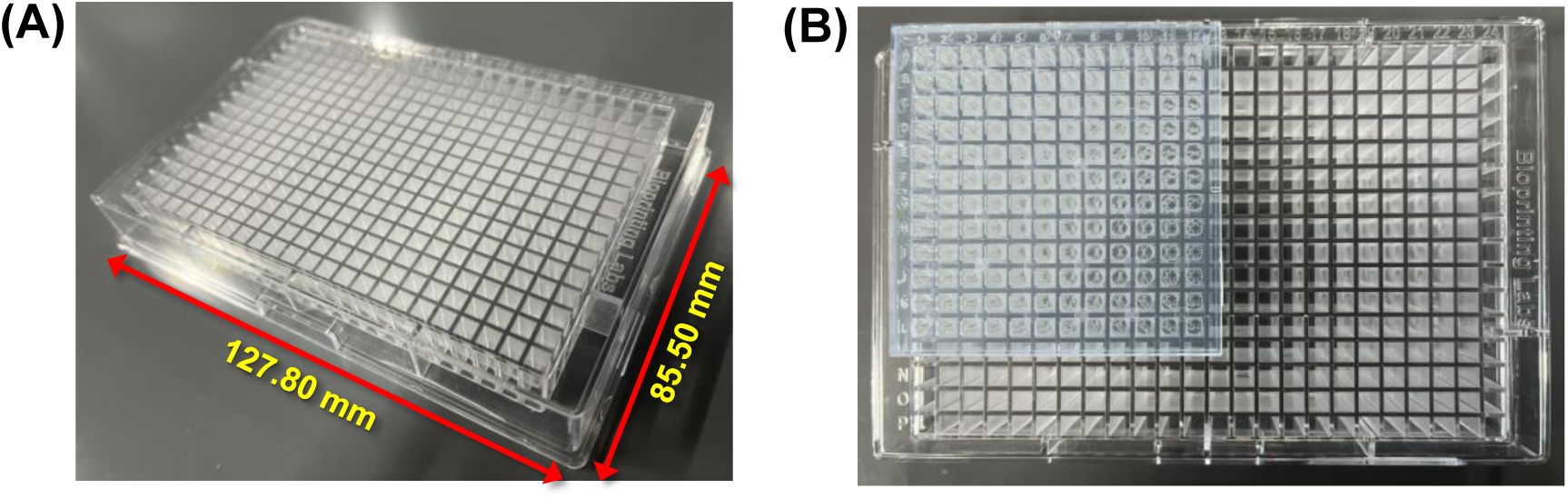
Injection-molded 384DeepWellPlate for static organoid culture on the 144PillarPlate: **(A)** The 384DeepWellPlate with a 16 x 24 array of deep wells. **(B)** The 144PillarPlate sandwiched onto the 384DeepWellPlate.

**Supplementary Figure 2.**
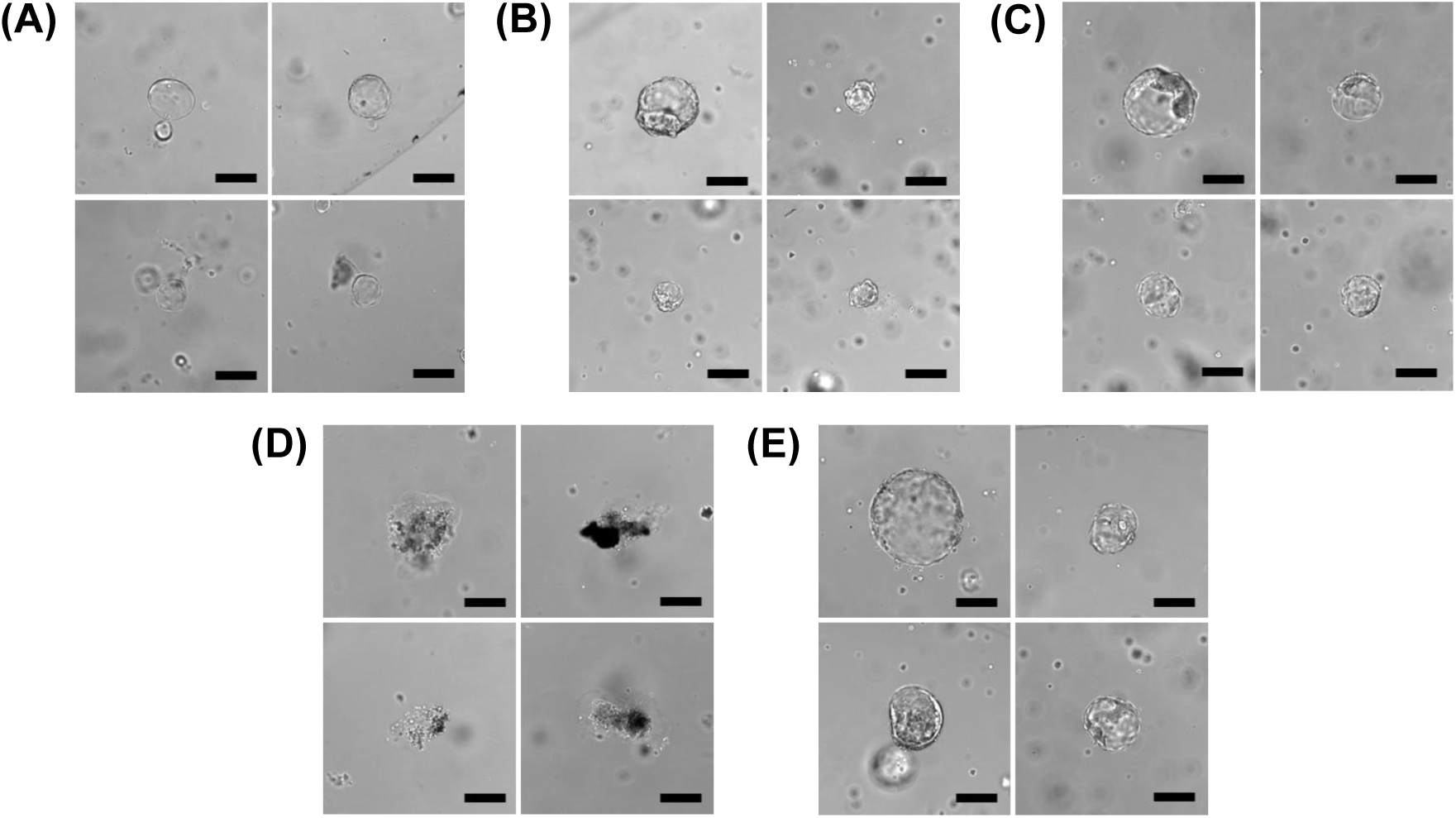
Optimization of the expansion medium (EM) to support culture of Exp- HLOs: **(A)** EM + Rspo1_HM + Wnt (initial 3 days) + Noggin (initial 3 days), **(B)** EM + Rspo1_HM, **(C)** EM + Rspo1_R&D, **(D)** EM - Rspo1 + Wnt + Noggin, and **(E)** EM - Rspo1 – EGF. Rspo1_HM and Rspo1_R&D represent home-made R-spondin1 conditioned medium and R-spondin1 purchased from R&D Systems, respectively. Rspo1_HM was prepared from the culture of Cultrex HA-R-Spondin1-Fc 293T cells (R&D Systems; 3710-001-01). The EM + Rspo1_HM + Wnt (3 days) + Noggin (3 days) represents the expansion medium supplemented with 10% (w/v) Rspo1_HM along with 30% (w/v) Wnt3a-conditioned medium (home-made) and 25 ng/mL Noggin for initial 3 days of culture. The EM + Rspo1_HM represents the expansion medium supplemented with 10% (w/v) home-made R-spondin1. The EM + Rspo1_R&D represents the expansion medium supplemented with 1 µg/mL R-spondin1 purchased from R&D Systems. The EM – Rspo1 + Wnt + Noggin represents the expansion medium supplemented with 30% (w/v) Wnt3a- conditioned medium (home-made) and 25 ng/mL Noggin without R-spondin1. The EM – Rspo1 – EGF represents the expansion medium without R-spondin1 and EGF. Scale bars: 200 μm.

**Supplementary Figure 3.**
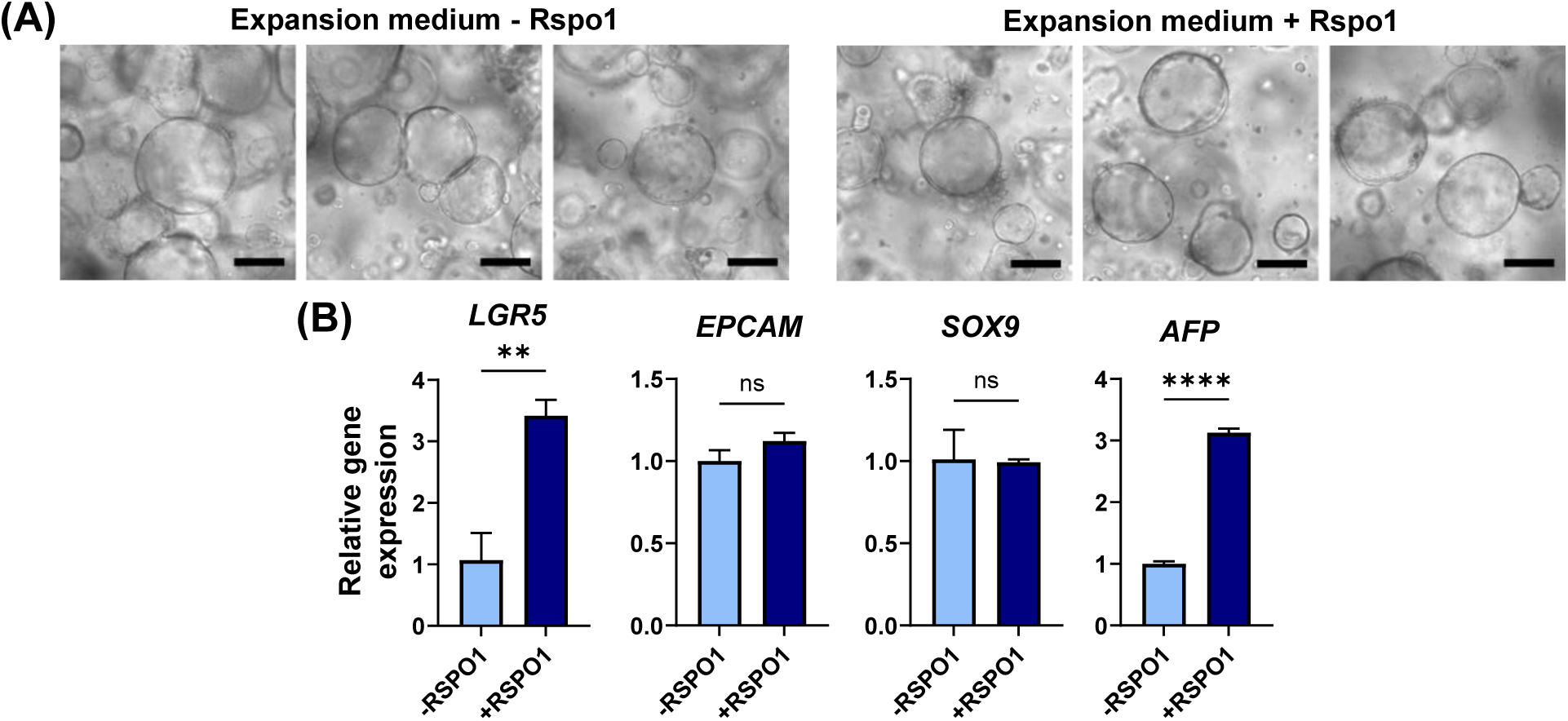
Optimization of the expansion medium (EM) to support culture of Exp- HLOs: **(A)** Representative images of Exp-HLOs cultured in the EM without and with R-spondin1 (Rspo1). Scale bars: 200 μm. **(B)** Comparison of hepatic gene expression in Exp-HLOs cultured in the EM without and with Rspo1, including *LGR5*, *EpCAM*, *SOX9*, and *AFP*. n = 3. The significance of gene expression difference was analyzed by Student’s t-test. Exp-HLOs at P4 were used for the differentiation into Diff- HLOs.

**Supplementary Figure 4.**
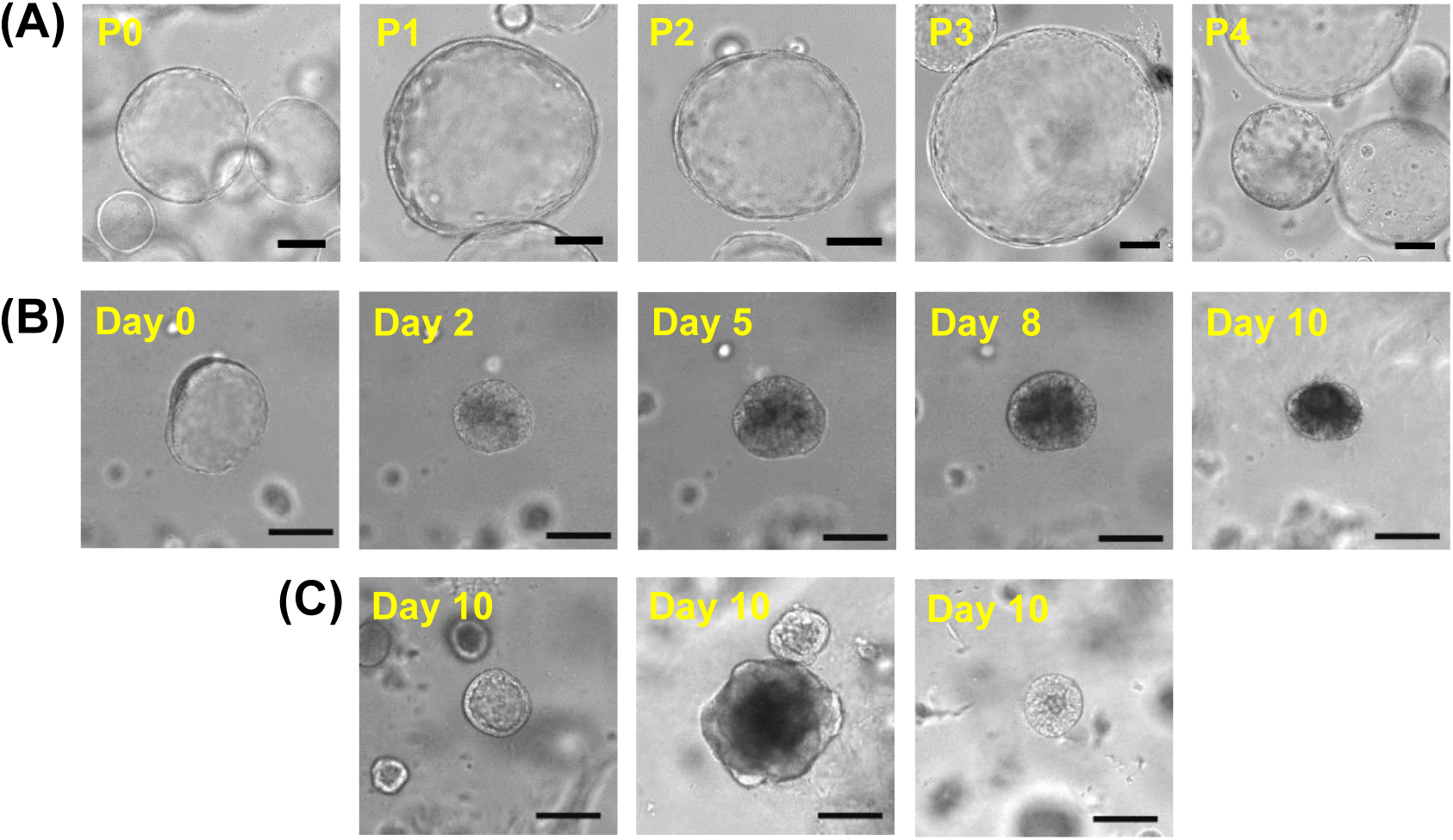
Passage of Exp-HLOs by mechanical dissociation *via* pipetting 10 - 15 times. **(A)** Representative images of passaged Exp-HLOs at different passage numbers. Scale bars: 200 μm. **(B)** Changes in morphology during differentiation of Exp-HLOs into Diff-HLOs for 10 days. Scale bars: 200 μm. **(C)** Representative images of day 10 Diff-HLOs. Scale bars: 200 μm. Exp-HLOs at P3 were used for the differentiation into Diff-HLOs.

**Supplementary Figure 5.**
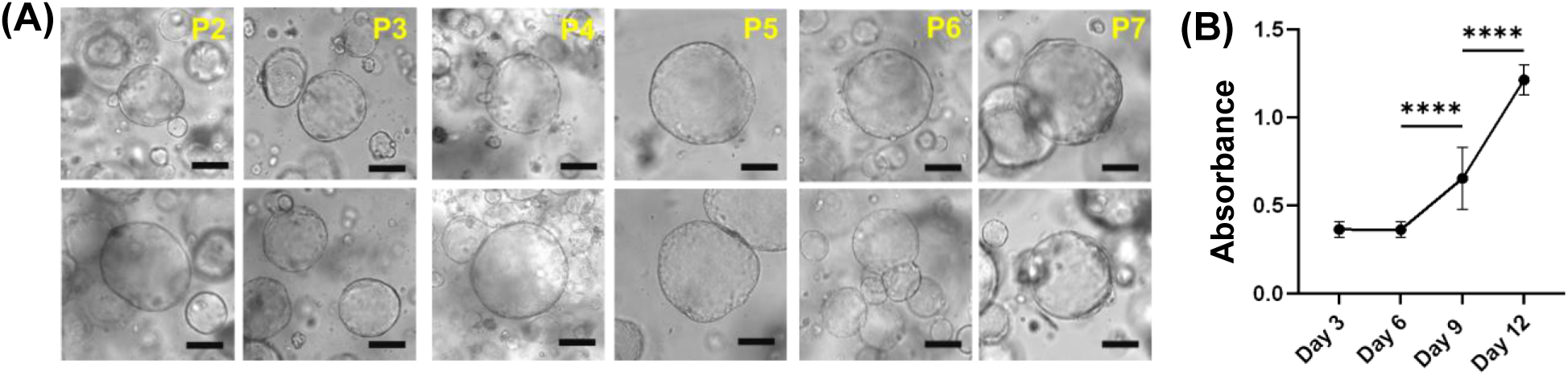
Cryopreservation of Exp-HLOs. **(A)** Morphology of Exp-HLOs before (P2, P4, and P6) and after (P3, P5, and P7) cryopreservation. Scale bars: 200 μm. **(B)** Proliferation of Exp-HLOs at P5 after 4 months of cryopreservation measured with the Cell Counting Kit 8. The significance was analyzed by Student’s t-test.

**Supplementary Figure 6.**
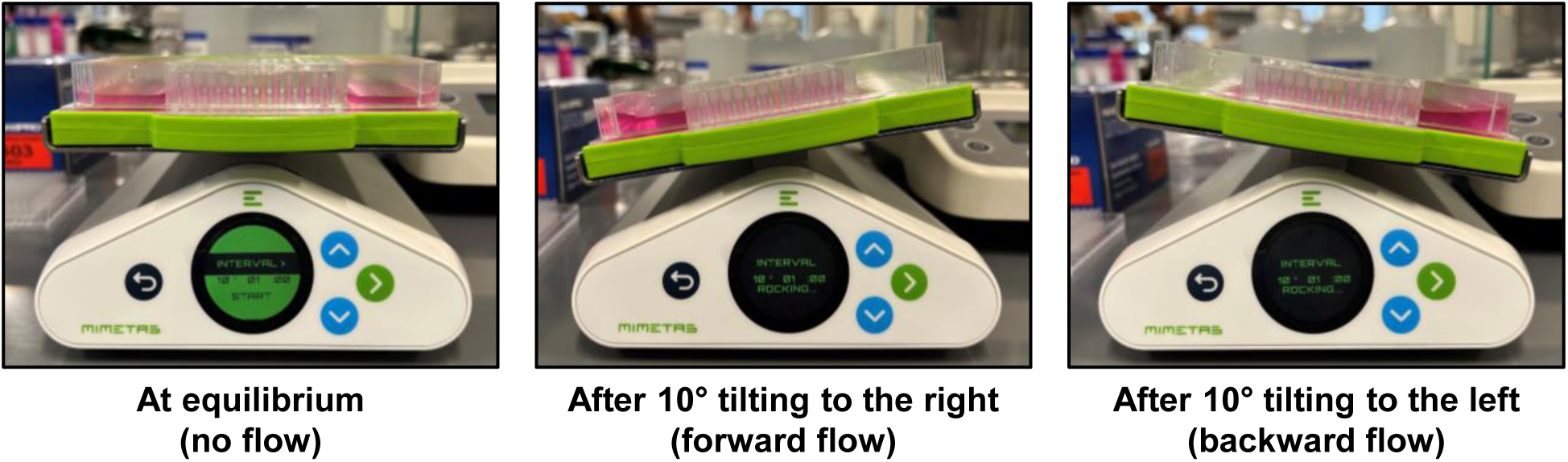
Dynamic culture of HLOs in the 144PillarPlate/144PerfusionPlate via gravity-driven, bi-directional flow on a digital rocker.

**Supplementary Figure 7.**
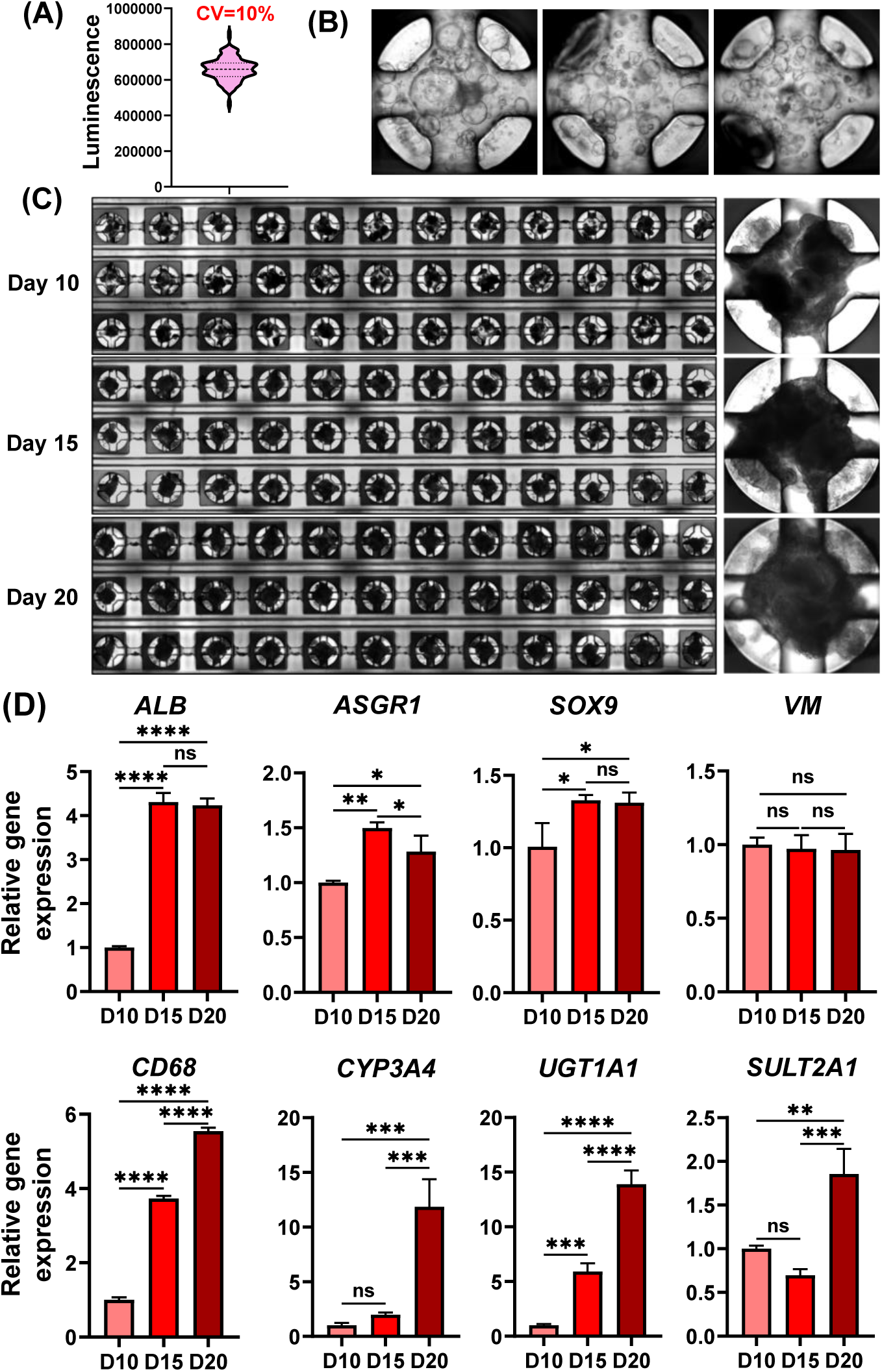
Bioprinting of Exp-HLOs on the pillar plate and dynamic differentiation into Diff-HLOs in the perfusion plate: **(A)** Uniformity of printing dissociated Exp-HLOs on the 144PillarPlate measured by using ATP-based cell viability assay kit. **(B)** Representative images of Exp- HLOs cultured on the 144PillarPlate. **(C)** Stitched (left) and representative (right) images of Diff-HLOs cultured in the 144PillarPlate/144PerfusionPlate for 10, 15, and 20 days. **(D)** Changes in hepatic gene expression in Diff-HLOs dynamically cultured for 10, 15, and 20 days, including *ALB*, *ASGR1*, *SOX9*, *VM*, *CD68*, *CYP3A4*, *UGT1A1*, and *SULT2A1*. n = 36. The significance of gene expression difference among the samples was analyzed using one-way ANOVA. Exp-HLOs at P6 were used for the differentiation into Diff-HLOs.

**Supplementary Figure 8.**
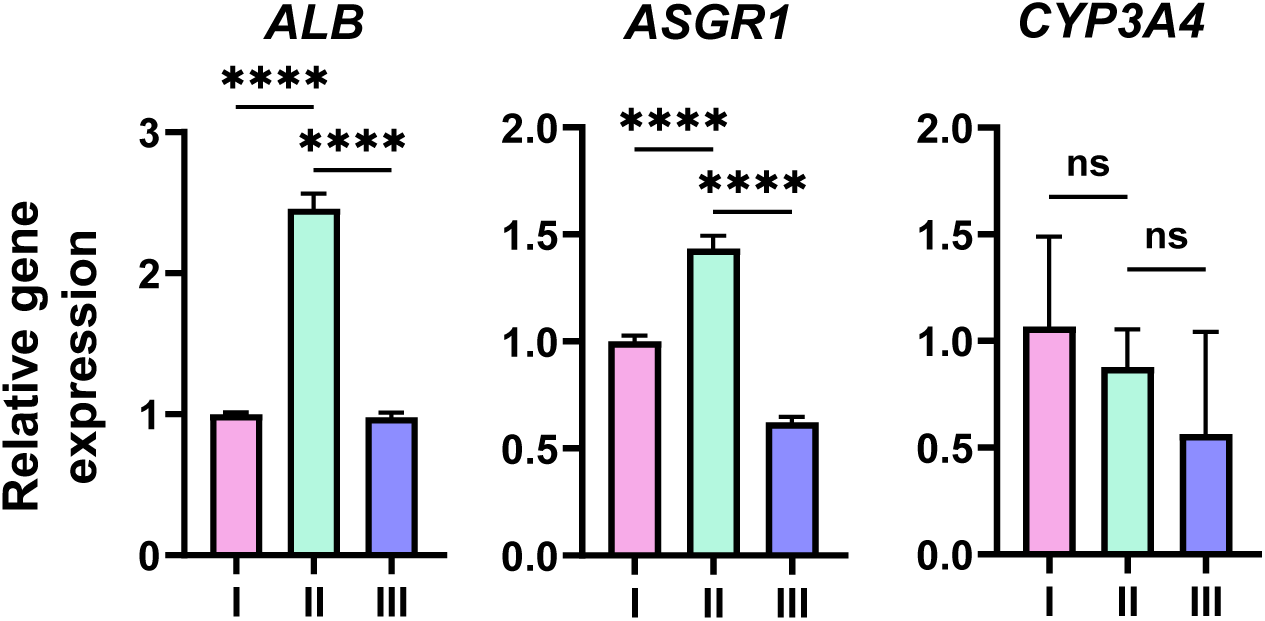
**Relative gene expression levels of *ALB*, *ASGR1*, and *CYP3A4*** compared among **(I)** Day 21, **(II)** Day 25, and **(III)** Day 30 HLOs cultured by using Ouchi *et al.* protocol. n = 3. Significant analysis was performed using one-way ANOVA.

**Supplementary Figure 9.**
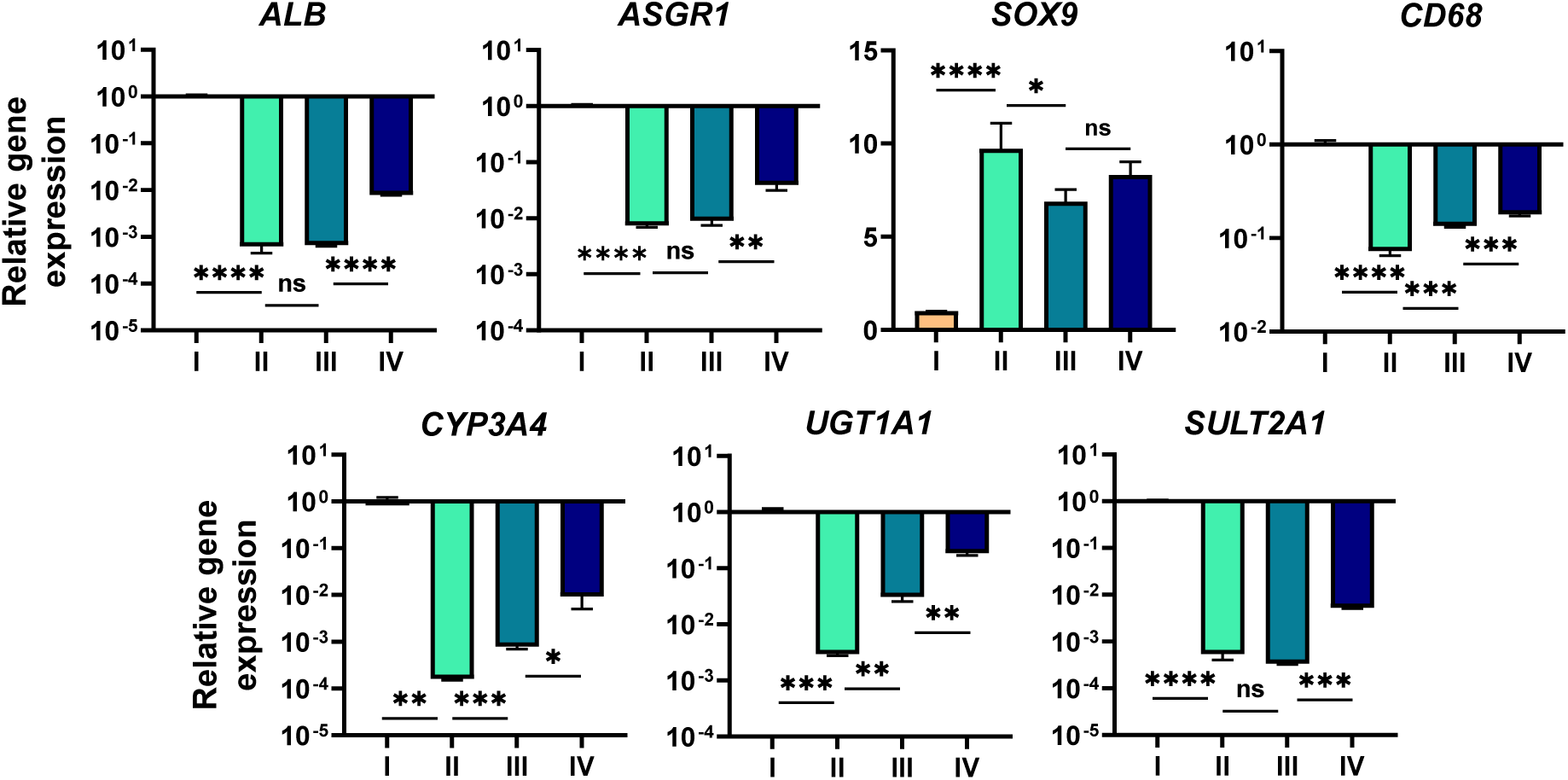
**Relative gene expression levels of *ALB*, *ASGR1*, *SOX9*, *CD68*, *CYP3A4*, *UGT1A1*, and *SULT2A1*** compared among **(I)** Human liver total RNA as a control (Takara; 636531), **(II)** Day 25 HLOs from Ouchi *et al*. protocol. n = 4, **(III)** Statically cultured Day 10 diff-HLOs in Matrigel dome. n = 4, and **(IV)** Dynamically cultured Day 20 diff-HLOs in the pillar/perfusion plate. n = 72. Significant analysis was performed using Student’s t-test.

## References

1 M. Gomez-Lechon, M. Donato, J. Castell and R. Jover, Curr Drug Metab, 2004, 5, 443–462.

2 S. J. Kwon, D. W. Lee, D. A. Shah, B. Ku, S. Y. Jeon, K. Solanki, J. D. Ryan, D. S. Clark, J. S. Dordick and M. Y. Lee, Nat Commun, 2014, 5, 3739.

3 A. Madan, R. A. Graham, K. M. Carroll, D. R. Mudra, L. A. Burton, L. A. Krueger, A. D. Downey, M. Czerwinski, J. Forster, M. D. Ribadeneira, L.-S. Gan, E. L. LeCluyse, K. Zech, P. Robertson, P. Koch, L. Antonian, G. Wagner, L. Yu and A. Parkinson, Drug Metabolism and Disposition, 2003, 31, 421–431.

4 H. H. J. Gerets, K. Tilmant, B. Gerin, H. Chanteux, B. O. Depelchin, S. Dhalluin and F. A. Atienzar, Cell Biol Toxicol, 2012, 28, 69–87.

5 D. T. U. H. Lam, Y. Y. Dan, Y.-S. Chan and H.-H. Ng, Cell Regeneration, 2021, 10, 27.

6 M. Huch, M. Grompe, A. Haft, H. Clevers, C. Dorrell, R. G. Vries, S. F. Boj, M. van de Wetering, T. Sato, N. Sasaki, M. J. Finegold, V. S. W. Li, K. Hamer and J. H. van Es, Nature, 2013, 494, 247– 250.

7 M. Huch, H. Gehart, R. Van Boxtel, K. Hamer, F. Blokzijl, M. M. A. Verstegen, E. Ellis, M. Van Wenum, S. A. Fuchs, J. De Ligt, M. Van De Wetering, N. Sasaki, S. J. Boers, H. Kemperman, J. De Jonge, J. N. M. Ijzermans, E. E. S. Nieuwenhuis, R. Hoekstra, S. Strom, R. R. G. Vries, L. J. W. Van Der Laan, E. Cuppen and H. Clevers, Cell, 2015, 160, 299–312.

8 H. Hu, H. Gehart, B. Artegiani, C. LÖ pez-Iglesias, F. Dekkers, O. Basak, J. van Es, S. M. Chuva de Sousa Lopes, H. Begthel, J. Korving, M. van den Born, C. Zou, C. Quirk, L. Chiriboga, C. M. Rice, S. Ma, A. Rios, P. J. Peters, Y. P. de Jong and H. Clevers, Cell, 2018, 175, 1591–1606.e19.

9 M. Chang, M. S. Bogacheva and Y.-R. Lou, Front Cell Dev Biol, DOI:10.3389/fcell.2021.748576.

10 M. Ogawa, S. Ogawa, C. E. Bear, S. Ahmadi, S. Chin, B. Li, M. Grompe, G. Keller, B. M. Kamath and A. Ghanekar, Nat Biotechnol, 2015, 33, 853–861.

11 F. Sampaziotis, M. Cardoso de Brito, P. Madrigal, A. Bertero, K. Saeb-Parsy, F. A. C. Soares, E. Schrumpf, E. Melum, T. H. Karlsen, J. A. Bradley, W. T. H. Gelson, S. Davies, A. Baker, A. Kaser, G. J. Alexander, N. R. F. Hannan and L. Vallier, Nat Biotechnol, 2015, 33, 845–852.

12 F. Wu, D. Wu, Y. Ren, Y. Huang, B. Feng, N. Zhao, T. Zhang, X. Chen, S. Chen and A. Xu, J Hepatol, 2019, 70, 1145–1158.

13 R. Ouchi, S. Togo, M. Kimura, T. Shinozawa, M. Koido, H. Koike, W. Thompson, R. A. Karns, C. N. Mayhew, P. S. McGrath, H. A. McCauley, R.-R. Zhang, K. Lewis, S. Hakozaki, A. Ferguson, N. Saiki, Y. Yoneyama, I. Takeuchi, Y. Mabuchi, C. Akazawa, H. Y. Yoshikawa, J. M. Wells and T. Takebe, Cell Metab, 2019, 30, 374–384.e6.

14 M. N. Bin Ramli, Y. S. Lim, C. T. Koe, D. Demircioglu, W. Tng, K. A. U. Gonzales, C. P. Tan, I. Szczerbinska, H. Liang, E. L. Soe, Z. Lu, C. Ariyachet, K. M. Yu, S. H. Koh, L. P. Yaw, N. H. B. Jumat, J. S. Y. Lim, G. Wright, A. Shabbir, Y. Y. Dan, H. H. Ng and Y. S. Chan, Gastroenterology, 2020, 159, 1471–1486.e12.

15 T. Shinozawa, M. Kimura, Y. Cai, N. Saiki, Y. Yoneyama, R. Ouchi, H. Koike, M. Maezawa, R.-R. Zhang, A. Dunn, A. Ferguson, S. Togo, K. Lewis, W. L. Thompson, A. Asai and T. Takebe, Gastroenterology, 2021, 160, 831–846.e10.

16 T. Takebe, K. Sekine, M. Enomura, H. Koike, M. Kimura, T. Ogaeri, R.-R. Zhang, Y. Ueno, Y.-W. Zheng, N. Koike, S. Aoyama, Y. Adachi and H. Taniguchi, Nature, 2013, 499, 481–484.

17 T. Takebe, K. Sekine, M. Kimura, E. Yoshizawa, S. Ayano, M. Koido, S. Funayama, N. Nakanishi, T. Hisai, T. Kobayashi, T. Kasai, R. Kitada, A. Mori, H. Ayabe, Y. Ejiri, N. Amimoto, Y. Yamazaki, S. Ogawa, M. Ishikawa, Y. Kiyota, Y. Sato, K. Nozawa, S. Okamoto, Y. Ueno and H. Taniguchi, Cell Rep, 2017, 21, 2661–2670.

18. A. Collin de l’Hortet, K. Takeishi, J. Guzman-Lepe, K. Morita, A. Achreja, B. Popovic, Y. Wang, K. Handa, A. Mittal, N. Meurs, Z. Zhu, F. Weinberg, M. Salomon, I. J. Fox, C.-X. Deng, D. Nagrath and A. Soto-Gutierrez, Cell Metab, 2019, 30, 385–401.e9.

19 S. Kang, M. Kimura, S. Shrestha, P. Lewis, S. Lee, Y. Cai, P. Joshi, P. Acharya, J. Liu, Y. Yang, J. G. Sanchez, S. Ayyagari, E. Alsberg, J. M. Wells, T. Takebe and M. Lee, Adv Healthc Mater, 2023, e2302502.

20 V. K. R. Lekkala, S.-Y. Kang, J. Liu, S. Shrestha, P. Acharya, P. Joshi, M. Zolfaghar, M. Lee, M. G. Vanga, P. Jamdagneya, S. Pagnis, A. Kundi, S. Kabbur, U. T. Kim, Y. Yang and M.-Y. Lee, ACS Biomater Sci Eng, 2024, 10, 3478–3488.

21. W. L. Thompson and T. Takebe, in Methods in Cell Biology, Academic Press Inc., 2020, vol. 159, pp. 47–68.

22 G. Chaturvedi, P. D. Simone, R. Ain, M. J. Soares and M. W. Wolfe, Cell Prolif, 2009, 42, 425– 433.

23 N. Lugli, I. Kamileri, A. Keogh, T. Malinka, M. E. Sarris, I. Talianidis, O. Schaad, D. Candinas, D. Stroka and T. D. Halazonetis, EMBO Rep, 2016, 17, 769–779.

24 G. K. Michalopoulos, W. C. Bowen, K. Mule and J. Luo, Gene Expr, 2018, 11, 55–75.

25 M. Kitade, V. M. Factor, J. B. Andersen, A. Tomokuni, K. Kaji, H. Akita, A. Holczbauer, D. Seo, J. U. Marquardt, E. A. Conner, S.-B. Lee, Y.-H. Lee and S. S. Thorgeirsson, Genes Dev, 2013, 27, 1706–1717.

26 H. E. Moad and A. A. Pioszak, Biochemistry, 2013, 52, 7295–7304.

27 A. Miyajima, T. Kinoshita, M. Tanaka, A. Kamiya, Y. Mukouyama and T. Hara, Cytokine Growth Factor Rev, 2000, 11, 177–183.

28 A. Kamiya, T. Kinoshita and A. Miyajima, FEBS Lett, 2001, 492, 90–94.

29 Acetaminophen, National Institute of Diabetes and Digestive and Kidney Diseases, Bethesda (MD), 2012.

30 M. C. Bouwmeester, Y. Tao, S. Proença, F. G. van Steenbeek, R.-A. Samsom, S. M. Nijmeijer, T. Sinnige, L. J. W. van der Laan, J. Legler, K. Schneeberger, N. I. Kramer and B. Spee, Molecules, 2023, 28, 621.

31 Troglitazone, National Institute of Diabetes and Digestive and Kidney Diseases, Bethesda (MD), 2012.

32 M. M. Clemens, M. R. McGill and U. Apte, 2019, pp. 241–262.

33 G. K. Michalopoulos and B. Bhushan, Nat Rev Gastroenterol Hepatol, 2021, 18, 40–55.

34 S. Hora and T. Wuestefeld, Cells, 2023, 12, 2129.

35 J. Qian, Y. Jiao, G. Wang, H. Liu, X. Cao and H. Yang, Exp Ther Med, 2020, 20, 1541–1549.

36 E. Luczak, J. Wieczfinska, M. Sokolowska, E. Pniewska, D. Luczynska and R. Pawliczak, Pharmacological Reports, 2017, 69, 1315–1321.

37 B. Bhushan and U. Apte, Am J Pathol, 2019, 189, 719–729.

38 S. J. Mun, J.-S. Ryu, M.-O. Lee, Y. S. Son, S. J. Oh, H.-S. Cho, M.-Y. Son, D.-S. Kim, S. J. Kim, H. J. Yoo, H.-J. Lee, J. Kim, C.-R. Jung, K.-S. Chung and M. J. Son, J Hepatol, 2019, 71, 970–985.

39 S. Wang, X. Wang, Z. Tan, Y. Su, J. Liu, M. Chang, F. Yan, J. Chen, T. Chen, C. Li, J. Hu and Y. Wang, Cell Res, 2019, 29, 1009–1026.

40 S. Akbari, G. G. Sevinç, N. Ersoy, O. Basak, K. Kaplan, K. Sevinç, E. Ozel, B. Sengun, E. Enustun, B. Ozcimen, A. Bagriyanik, N. Arslan, T. T. Ö nder and E. Erdal, Stem Cell Reports, 2019, 13, 627– 641.

41 H. Kim, I. Im, J. S. Jeon, E.-H. Kang, H.-A. Lee, S. Jo, J.-W. Kim, D.-H. Woo, Y. J. Choi, H. J. Kim, J.-S. Han, B.-S. Lee, J.-H. Kim, S. K. Kim and H.-J. Park, Biomaterials, 2022, 286, 121575.

42 M. Hofer and M. P. Lutolf, Nat Rev Mater, 2021, 6, 402–420.

43 Y. Wang, L. Wang, Y. Guo, Y. Zhu and J. Qin, RSC Adv, 2018, 8, 1677–1685.

44 M. J. Workman, J. P. Gleeson, E. J. Troisi, H. Q. Estrada, S. J. Kerns, C. D. Hinojosa, G. A. Hamilton, S. R. Targan, C. N. Svendsen and R. J. Barrett, CMGH, 2018, 5, 669–677.e2.

45 Y. Wang, H. Wang, P. Deng, W. Chen, Y. Guo, T. Tao and J. Qin, Lab Chip, 2018, 18, 3606–3616.

46 E. Berger, C. Magliaro, N. Paczia, A. S. Monzel, P. Antony, C. L. Linster, S. Bolognin, A. Ahluwalia and J. C. Schwamborn, Lab Chip, 2018, 18, 3172–3183.

47 K. A. Homan, N. Gupta, K. T. Kroll, D. B. Kolesky, M. Skylar-Scott, T. Miyoshi, D. Mau, M. T. Valerius, T. Ferrante, J. V. Bonventre, J. A. Lewis and R. Morizane, Nat Methods, 2019, 16, 255– 262.

48 T. Tao, Y. Wang, W. Chen, Z. Li, W. Su, Y. Guo, P. Deng and J. Qin, Lab Chip, 2019, 19, 948– 958.

49 A. N. Cho, Y. Jin, Y. An, J. Kim, Y. S. Choi, J. S. Lee, J. Kim, W. Y. Choi, D. J. Koo, W. Yu, G. E. Chang, D. Y. Kim, S. H. Jo, J. Kim, S. Y. Kim, Y. G. Kim, J. Y. Kim, N. Choi, E. Cheong, Y. J. Kim, H. S. Je, H. C. Kang and S. W. Cho, Nat Commun, DOI:10.1038/s41467-021-24775-5.

50 P. Saglam-Metiner, U. Devamoglu, Y. Filiz, S. Akbari, G. Beceren, B. Goker, B. Yaldiz, S. Yanasik, C. Biray Avci, E. Erdal and O. Yesil-Celiktas, Commun Biol, DOI:10.1038/s42003-023-04547-1.

51 T. Mammoto and D. E. Ingber, Development, 2010, 137, 1407–1420.

52 S. Vianello and M. P. Lutolf, Dev Cell, 2019, 48, 751–763.

53 J. Sia, R. Sun, J. Chu and S. Li, Biomaterials, 2016, 92, 36–45.

54 L. Ewart, A. Apostolou, S. A. Briggs, C. V. Carman, J. T. Chaff, A. R. Heng, S. Jadalannagari, J. Janardhanan, K.-J. Jang, S. R. Joshipura, M. M. Kadam, M. Kanellias, V. J. Kujala, G. Kulkarni, C. Y. Le, C. Lucchesi, D. V. Manatakis, K. K. Maniar, M. E. Quinn, J. S. Ravan, A. C. Rizos, J. F. K. Sauld, J. D. Sliz, W. Tien-Street, D. R. Trinidad, J. Velez, M. Wendell, O. Irrechukwu, P. K. Mahalingaiah, D. E. Ingber, J. W. Scannell and D. Levner, Communications Medicine, 2022, 2, 154.

55 S. A. P. Rajan, J. Sherfey, S. Ohri, L. Nichols, J. T. Smith, P. Parekh, E. P. Kadar, F. Clark, B. T. George, L. Gregory, D. Tess, J. R. Gosset, J. Liras, E. Geishecker, R. S. Obach and M. Cirit, AAPS J, 2023, 25, 102.

56 L. Docci, N. Milani, T. Ramp, A. A. Romeo, P. Godoy, D. O. Franyuti, S. Krähenbühl, M. Gertz, A. Galetin, N. Parrott and S. Fowler, Lab Chip, 2022, 22, 1187–1205.

57 S.-Y. Chang, J. L. Voellinger, K. P. Van Ness, B. Chapron, R. M. Shaffer, T. Neumann, C. C. White, T. J. Kavanagh, E. J. Kelly and D. L. Eaton, Toxicology in Vitro, 2017, 40, 170–183.

58 A. Iqbal, N. Van Hul, L. Belicova, A. A. Corbat, S. Hankeova and E. R. Andersson, Liver International, 2024, 44, 541–558.

59 S. Shrestha, V. K. R. Lekkala, P. Acharya, S.-Y. Kang, M. G. Vanga and M.-Y. Lee, Lab Chip, 2024, 24, 2747–2761.

60 P. Acharya, P. Joshi, S. Shrestha, N. Y. Choi, S. Jeong and M.-Y. Lee, Biofabrication, 2024, 16, 025005.

61 P. Acharya, S. Shrestha, P. Joshi, N. Y. Choi, V. K. R. Lekkala, S.-Y. Kang, G. Ni and M.-Y. Lee, Biofabrication, 2025, 17, 015001.

62 R. S. Mangipudy, S. Chanda and H. M. Mehendale, Environ Health Perspect, 1995, 103, 260–267.

63 B. Bhushan, C. Walesky, M. Manley, T. Gallagher, P. Borude, G. Edwards, S. P. S. Monga and U. Apte, Am J Pathol, 2014, 184, 3013–3025.

64 T. G. Bird, M. Müller, L. Boulter, D. F. Vincent, R. A. Ridgway, E. Lopez-Guadamillas, W.-Y. Lu, T. Jamieson, O. Govaere, A. D. Campbell, S. Ferreira-Gonzalez, A. M. Cole, T. Hay, K. J. Simpson, W. Clark, A. Hedley, M. Clarke, P. Gentaz, C. Nixon, S. Bryce, C. Kiourtis, J. Sprangers, R. J. B. Nibbs, N. Van Rooijen, L. Bartholin, S. R. McGreal, U. Apte, S. T. Barry, J. P. Iredale, A. R. Clarke, M. Serrano, T. A. Roskams, O. J. Sansom and S. J. Forbes, Sci Transl Med, DOI:10.1126/scitranslmed.aan1230.

65 S. J. Mun, Y.-H. Hong, Y. Shin, J. Lee, H.-S. Cho, D.-S. Kim, K.-S. Chung and M. J. Son, Sci Rep, 2023, 13, 22935.

66 S. V. Beath, Seminars in Neonatology, 2003, 8, 337–346.

67 J. J. Velazquez, R. LeGraw, F. Moghadam, Y. Tan, J. Kilbourne, J. C. Maggiore, J. Hislop, S. Liu, D. Cats, S. M. Chuva de Sousa Lopes, C. Plaisier, P. Cahan, S. Kiani and M. R. Ebrahimkhani, Cell Syst, 2021, 12, 41–55.e11.

